# Healthy aging delays and dedifferentiates high-level visual representations

**DOI:** 10.1101/2024.07.30.605732

**Authors:** Marleen Haupt, Douglas D. Garrett, Radoslaw M. Cichy

## Abstract

Healthy aging impacts visual information processing with consequences for subsequent high-level cognition and everyday behavior, but the underlying neural changes in visual representations remain unknown. Here, we investigate the nature of representations underlying object recognition in older compared to younger adults by tracking them in time using EEG, across space using fMRI, and by probing their behavioral relevance using similarity judgements. Applying a multivariate analysis framework to combine experimental assessments, four key findings about how brain aging impacts object recognition emerge. First, aging selectively delays the formation of object representations, profoundly changing the chronometry of visual processing. Second, the delay in the formation of object representations emerges in high-level rather than low- and mid-level ventral visual cortex, supporting the theory that brain areas developing last deteriorate first. Third, aging reduces content selectivity in high-level ventral visual cortex, indicating age-related neural dedifferentiation as the mechanism of representational change. Finally, we demonstrate that the identified representations of the aging brain are behaviorally relevant, ascertaining ecological relevance. Together, our results reveal the impact of healthy aging on the visual brain.

## 2. Introduction

Aging is associated with cognitive declines impacting older adults’ everyday competences. Even in the absence of pathological brain changes, healthy aging decreases processing speed, executive functions as well as episodic and working memory performance ^2,3^. While high-level cognitive functions are often the focus of investigation and intervention in older adults, aging already profoundly impairs the preceding stages of visual cortical information processing ^4^.

Importantly, changes in visual cortical information processing alter how older participants perceive the world, including, for example, how they detect motions, identify directions, and perceive speed ^4–6^ in everyday situations such as driving. Differences at this core visual processing level in turn impact high-level cognitive performance in older adults, both in experimental assessment that is often visual and in everyday life. Thus, quantifying age-related differences in basic visual information processing is integral to studying cognitive aging.

Previous aging studies have shown prolonged latencies, decreased amplitudes and reduced distinctiveness of brain responses to visual stimulation ^7–9^. While these findings firmly establish an impact of healthy aging on visual processing, they do not yet provide insight into how the underlying neural representations themselves are affected to cause the observed changes. In this study, we aim to reveal the nature of representations underlying visual object processing in older adults to characterize how brain aging impacts the neural representations underlying object recognition.

In order to provide a comprehensive view on object recognition in healthy aging, we measure both EEG and fMRI responses to natural images and behaviorally assess the perceptual similarity of these stimuli in younger and older participants. Combining the neural measurements with multivariate analysis techniques ^10–13^, we determine the chronometry underlying the formation of object representations, localize them to cortical regions, and characterize the representations there. We finally ascertain the ecological relevance of the identified neural representations by relating them to the behavioral assessment of perceived similarity ^14–17^.

## 3. Results

We conducted three separate experimental sessions in younger (19-33 years) and cognitively healthy, older participants (60-73 years) (for details see Table 1): two fMRI sessions and one session separately assessing EEG and behavior. We recorded EEG and fMRI separately while participants viewed natural images. The experimental design was identical across age groups to enable robust comparisons, and the stimulus set was comparable for both imaging modalities to enable easy integration. The stimulus set consisted of 64 images belonging to the four categories faces, animals, places and objects (i.e., 16 per category, see Fig. 1a). Five different images of smurfs in the EEG paradigm and a fixation cross color change in the fMRI paradigm served as vigilance targets. We instructed participants to respond to catch trials with a button press and excluded catch trials from all analyses. In EEG, we presented images for 500ms with a jittered ISI of 500ms in target trials or 1500ms in catch trials (Fig. 1b left). In fMRI, we presented stimuli in a 750ms on-off-on-off design (Fig. 1b right) ^18^. We also assessed visual behavior through a multiple arrangement task, requiring participants to arrange the same 64 stimuli according to their perceived similarity.

We then used a set of multivariate pattern analysis tools to comprehensively characterize the nature of object representations in healthy aging, assessing representations in their temporal, spatial, and functional aspects.

**Table 1.** Participant information. MMSE: Mini-Mental State Examination, maximum score 30 points with values < 27 points indicating mild cognitive impairment.

|  | Younger adults<br>n = 21 | Older adults<br>n = 21 |
| --- | --- | --- |
| Age | 24.8 (4.8) | 66.7 (4.2) |
| Gender | 10 F / 11 M | 11 F / 10 M |
| Education (school years) | 12.4 (0.9) | 11.2 (1.3) |
| Cognition (MMSE) | - | 29.1 (0.8) |

**Figure 1.**
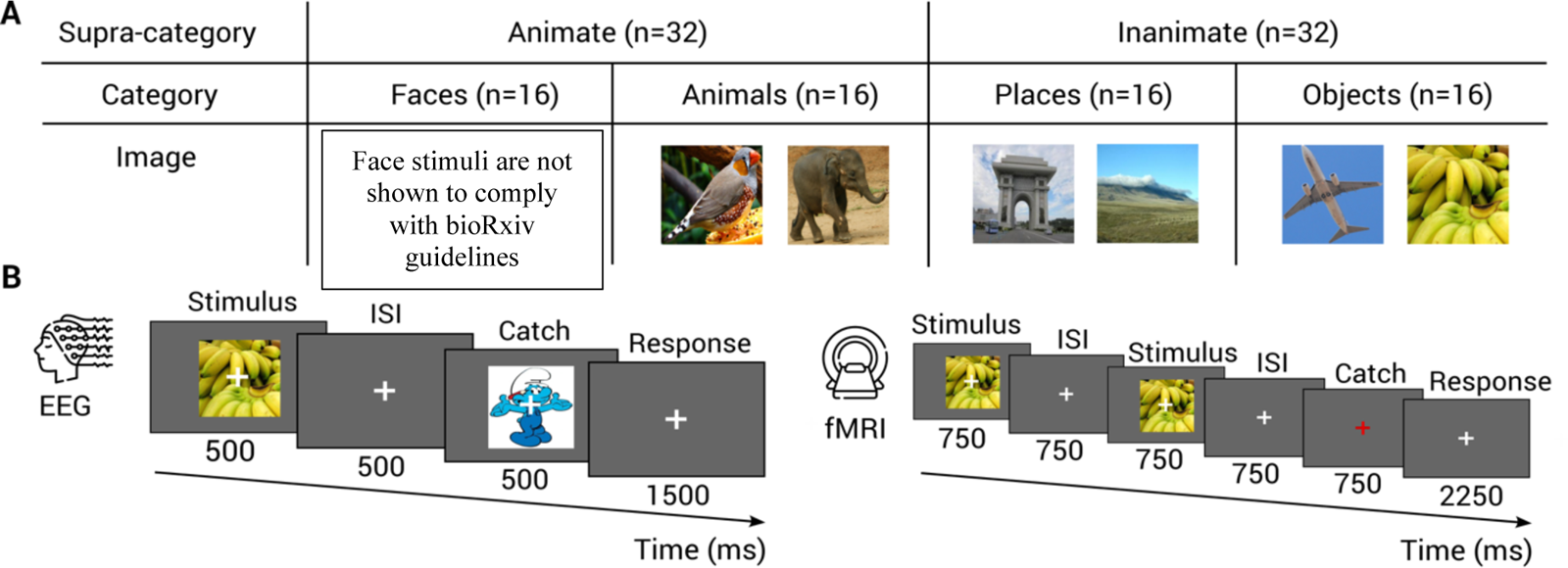
Stimuli and experimental design. A) The stimulus set consisted of 64 natural images from the categories faces, animals, places and objects. B) In both neuroimaging experiments, we presented stimuli on a grey background, overlayed with a central fixation cross. We instructed participants to fixate centrally and respond to catch trials by pressing a button. Left: During the EEG session, stimuli were presented for 500ms. The ISI was jittered around 500ms following regular trails and 1,500ms following catch trials to avoid movement confounds. Catch trials consisted of smurf images. Right: During a separate fMRI session, stimuli were presented in an 750ms on-off-on-off design. Catch trials consisted of fixation cross color changes.

### 3.1 The temporal dynamics of object representations

**Figure 2.**
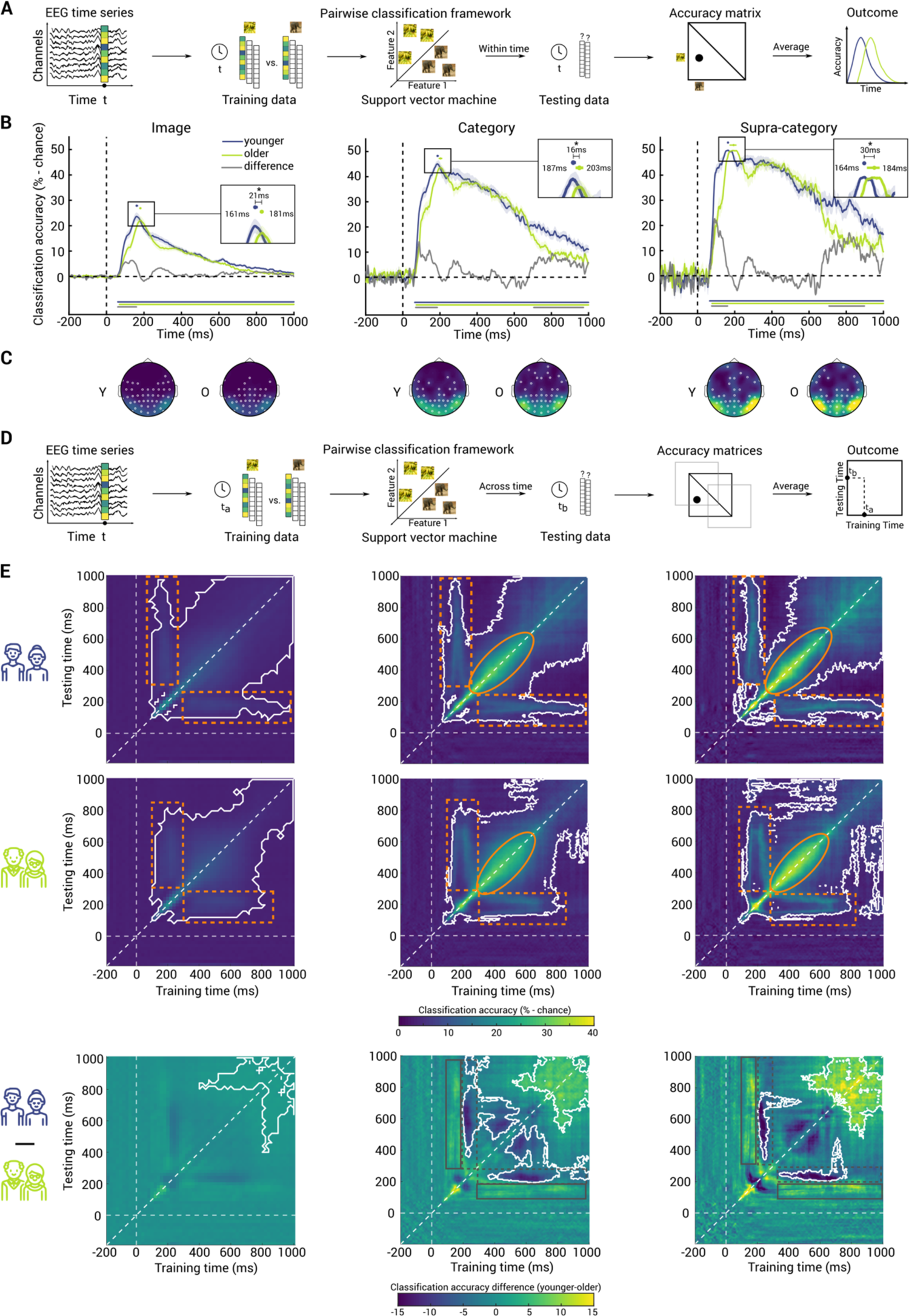
EEG classification within and across time. A) We divided EEG pattern vectors into training and test sets. For every pair of stimuli, we trained a support vector machine (SVM) to classify between pattern vectors related to the presentation of two images from the same time point. We tested the SVM on the left-out pattern vectors for the same images from the same time point. We aggregated the resulting pairwise classification accuracies in a matrix that is symmetric along the diagonal, with the diagonal itself being undefined. After averaging all pairwise classification accuracies, we plot accuracy over time. B) EEG time classification for image (left), category (middle) and supra-category (right) in younger adults (blue), older adults (green) and younger-older (grey). Shaded areas around curves indicate standard error of the mean. Significance time points are indicated below curves (10,000 permutations, one-tailed permutation test, p<0.05, cluster threshold q<0.05). Dots above curves indicate peaks of the classification curves and error bars represent the 95% CI. Stars indicate significant peak latency differences between age groups (p < 0.05; bootstrap test with 10,000 bootstraps). Insets indicate peak latency for younger adults on the left, peak latency for older adults on the right and peak latency differences on the top. Stars indicate significant peak latency differences between age groups (p < 0.05; bootstrap test with 10,000 bootstraps). C) Brains depict EEG searchlight classification results for younger (left) and older (right) adults at classification peaks in EEG channel space. Significant electrodes are marked with white circles (10,000 permutations, one-tailed permutation test, p<0.05, FDR corrected). D) In the case of time generalization, we train the SVM on vectors from time point t_a_ and test it on vectors from all other time points t_b_. We get one classification accuracy matrix per time combination and a time-time matrix after averaging across pair-wise entries. E) EEG time generalization for image (left), category (middle) and supra-category (right) in younger adults (top), older adults (middle) and age group difference (bottom). Dashed white lines indicate stimulus onset and the diagonal. Solid white outlines indicate significance clusters (10,000 permutations, one-tailed permutation test, p<0.05, cluster threshold q<0.05). Striped, orange rectangles indicate vertical and horizontal bands of persistent activity, orange ovals indicate a widening of the diagonal. In the age group comparison, solid grey rectangles indicate higher bands in younger adults and striped, grey rectangles indicate higher bands in older adults.

#### 3.1.1 Delayed visual processing in older adults

We commenced the characterization of object representations by assessing the temporal dynamics with which representations emerge in healthy older compared to younger adults. In a first step, we used time-resolved classification from EEG electrode patterns to determine when object information emerges at three levels of abstraction: image, category (faces, animals, places, objects) and supra-category (animate, inanimate) (Fig. 2a). We evaluated the classification time courses statistically using non-parametric permutation tests for one sample when assessing age groups separately and for independent samples when assessing age group differences. We controlled for multiple comparisons using cluster-based correction (10,000 permutations, one-tailed permutation test, p<0.05, cluster threshold q<0.05).

As expected, we found significant classification for both age groups (younger: blue curve and older: green curve) and at every abstraction level (Fig. 2b) with a stereotypical shape: classification curves rose steeply around 60ms and after a peak decayed gradually.

In a next step, we compared classification accuracies for younger and older adults in two ways: comparison of peak latency and inspection of the difference curve. Comparing peak latencies of the EEG classification time courses (for details including 95% confidence intervals see Supplementary Table 1) was motivated by the idea that classification peaks indicate when object representations are most distinct in terms of linear separability ^19^. We found that classification curves in older participants consistently peaked around 16-30ms later than in younger participants (Fig. 2b insets; 10,000 bootstraps, bootstrap test against zero, p<0.05). This indicates that processing in older participants is delayed compared to younger participants.

We then compared the results curves of younger and older participants (Fig. 2b, gray curves indicate difference younger-older). We found higher classification for younger participants at all three abstraction levels in an early time window around 60-160ms. This time frame overlaps with the timing of the feed-forward sweep across the ventral visual cortex ^19,20^ indicating that the observed delay in the older participants is due to delayed feed-forward processing. Notably, classification results for younger and older adults did not differ significantly at the time of peak classification, suggesting that potential differences between age groups are not trivially due to differences in signal-to-noise ratio. This establishes the feasibility to assess and compare visual object representations in both age groups and across stimulus abstraction levels.

Additionally, at the level of category and supra-category classification, younger adults showed higher classification in a late time window around 700-900ms. These late processing differences are beyond the timing of stimulus presentation and core visual processing and might be explained by higher sustained attention in younger adults ^21,22^.

To approximate the sources of the observed classification results, we conducted a searchlight classification analysis in EEG sensor space at peak latency. As expected, for both groups and at all abstraction levels, classification was strongest from occipital and temporal electrodes (Fig. 2c; 10,000 permutations, one-tailed permutation test, FDR corrected, p<0.05). This suggests similar cortical loci of the identified visual representations in the brains of younger and older adults.

#### 3.1.2 Comparable persistent and transient dynamics across age groups

We further characterized the neural dynamics in younger and older adults by disentangling the contributions of transient versus persistent neural components to the time-resolved results above. To achieve this, we used time generalization analysis (TGA, Fig. 2d) ^23,24^, classification across all time-point combinations in the epoch. This resulted in 2D matrices of classification accuracies indexed in rows and columns by the time points compared. High classification accuracies on the diagonal indicate rapidly changing transient representations while high classification accuracies off the diagonal indicate persistent representations that are maintained longer. As before, we assessed significance using one sample (younger and older adults separately) and independent samples (age group comparison) permutation tests with cluster-based correction (10,000 permutations, one-tailed permutation test, p<0.05, cluster threshold q<0.05).

Consistent with the overall similarity of the time-resolved classification curves across age groups, we observed similar TGA result patterns across age groups and at all abstraction levels (Fig. 2e top and middle row). In each case, we found a stereotypical picture observed previously ^23,24^ with a mixture of transient and persistent aspects. Strong transient dynamics were indicated by consistently highest classification accuracy on and close to the diagonal. Persistent dynamics were indicated by vertical and horizontal bands (striped orange rectangles), a widening of the diagonal across time (solid orange ellipse), and plateaus of classification at late processing stages.

A direct comparison of TGA results across age groups revealed two key differences (Fig. 2e bottom row). First, we observed bands of higher classification for older participants at the level of both category and supra-category (gray striped rectangles). Given that those bands follow bands of higher classification for younger participants (albeit effects are not significant, gray solid rectangles), and the timing corresponds to the respective peak latencies in the time-resolved classification, this indicates a delay in which persistent representations emerge in older participants. Second, we observed a highly persistent pattern at late stages of processing after 700ms, clarifying that the late effects observed in time-resolved classification reflect persistent representations.

Together, our results reveal that transient and persistent dynamics contribute similarly to visual object processing in younger and older adults. Yet, older participants exhibit delayed early persistent dynamics, while younger participants show additional late persistent dynamics.

#### 3.1.3 Younger and older adults rely on similar, time-shifted representations

**Figure 3.**
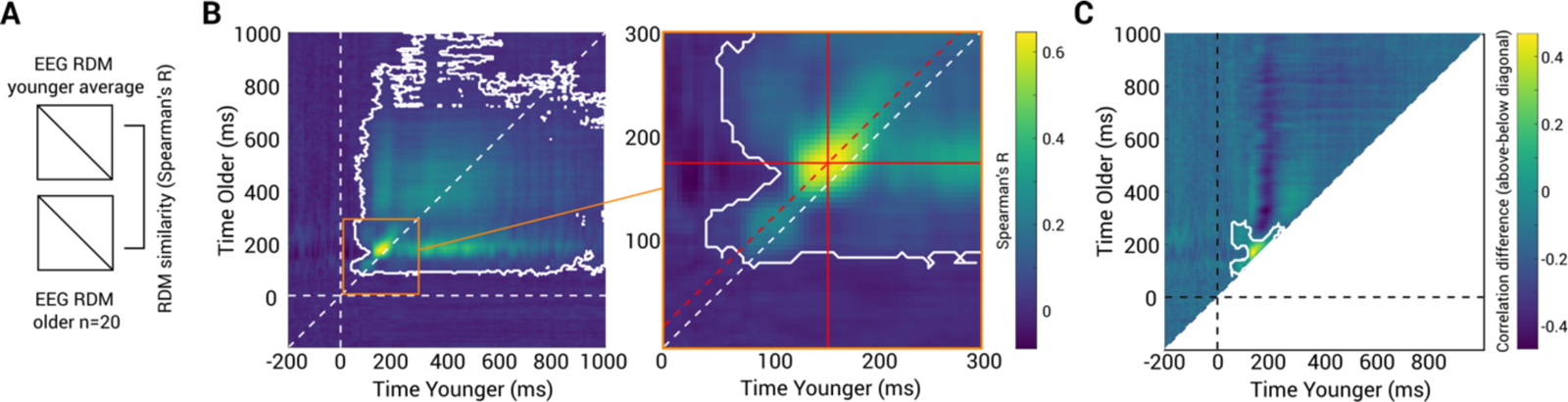
Comparing EEG RDMs across age groups. A) We correlated the individual EEG RDMs of older participants with the average EEG RDM of younger adults using Spearman’s R. B) Time-point wise correlations of older participants’ EEG RDMs with average younger EEG RDMs. Dashed white line indicates diagonal; solid white outlines indicate significant clusters of higher correlations above as compared to below the diagonal (10,000 permutations, one-tailed permutation test, p<0.05, cluster threshold q<0.05). Solid red lines indicate maximum correlation. Dashed red line indicates the offset from the diagonal for the maximum correlation value. D) Time-point wise correlations of older participants’ EEG RDMs with average younger EEG RDMs displayed as difference scores (above diagonal minus below diagonal). Solid white outlines indicate significant clusters of higher correlations above as compared to below the diagonal (10,000 permutations, one-tailed permutation test, p<0.05, cluster threshold q<0.05).

The temporal analyses so far showed similar but delayed results patterns for older compared to younger adults. This suggests that both age groups rely on similar, yet time-shifted visual representations. To directly test this hypothesis, we compared representations across age groups using representational similarity analysis (RSA) ^25^. We used the time-resolved image classification results as representational dissimilarity matrices ^24,26^, summarizing the representational geometry at each time point. We then correlated the individual time-resolved RDMs of older adults to the average time-resolved RDM of younger adults for all time-point combinations (Fig. 3a). The resulting 2D matrix, indexed by time in younger on the x-axis and time in older on the y-axis, showed significantly correlated representations emerging at around 100ms and lasting throughout the entire epoch (Fig. 3b; 10,000 permutations, one-tailed permutation test, p<0.05, cluster threshold q<0.05). We observed highest correlations above the diagonal, indicating that earlier time points in younger participants are correlated with later time points in older participants. We ascertained this qualitative observation in two ways. First, we determined the location of the peak correlation values by calculating their distance to the diagonal. We found that the maximum correlation values were located above the diagonal (p=0.0001, 10,000 permutations, one-tailed permutation test, p<0.05, FDR corrected). Second, we subtracted the lower triangular part of the matrix from the upper part to directly compare classification accuracy above and below the diagonal. This off-diagonal correlation difference was significant at around 50-250ms (Fig. 3c; 10,000 permutations, one-tailed permutation test, p<0.05, cluster threshold q<0.05). Together, these findings confirm the hypothesis that object representations in older adults are similar to those in younger adults but shifted in time.

#### 3.1.4 Robust age group classification depends on temporal processing delays

**Figure 4.**
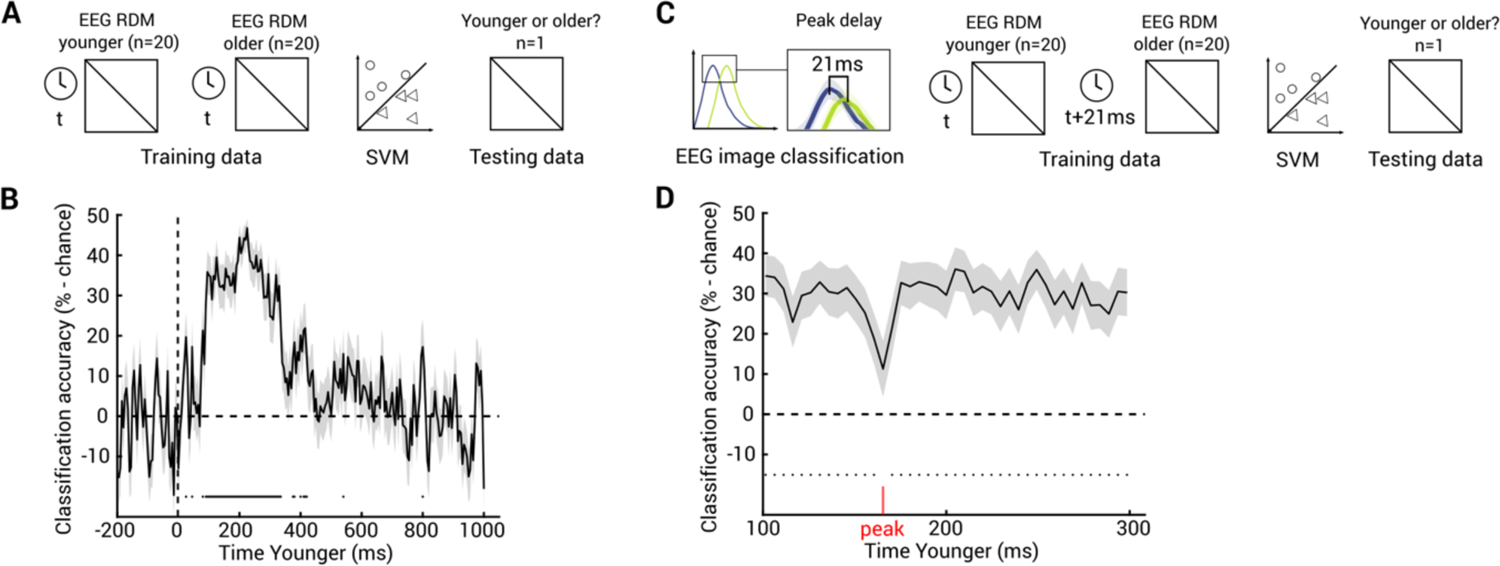
Age group classification from EEG RDMs. A) We trained a support vector machine (SVM) to classify between the EEG RDMs of 20 younger and 20 older participants and tested in on the RDM of a left-out participant, using the same time points for younger and older adults. B) Time course of age group classification based on EEG RDMs from the same time points. Shaded areas around curves indicate standard error of the mean. Significance time points are indicated below curves (10,000 permutations, one-tailed permutation test, p<0.05, FDR corrected). C) We used the same leave-one-participant-out procedure as in A but accounted for the age-related time shift by using time points t for younger adults and time points t+21ms (image classification delay) for older adults. D) Time course of age group classification based on EEG RDMs and accounting for peak delay in older. Importantly, x-axis indicates time points in younger adults. Shaded areas around curves indicate standard error of the mean. Significance time points are indicated below curves (10,000 permutations, one-tailed permutation test, p<0.05, FDR corrected). Red line indicates peak of image classification in younger adults.

The identified differences in the temporal dynamics between older and younger adults suggest two predictions that, if born out, would strengthen the results: first, that age groups are differentiable by their temporal dynamics, and second, that this differentiation depends on processing delays.

We tested these predictions by classifying age group based on time-resolved EEG RDMs, training classifiers on RDMs from younger versus older participants and testing them on a left-out participant. To test the first prediction, we conducted the analysis in a time-resolved fashion using the same time points for training and testing (Fig. 4a). The result confirmed the first prediction, showing age group classification based on participants’ EEG RDMs in a continuous time window from 90-335ms (Fig. 4b; 10,000 permutations, one-tailed permutation test, p<0.05, FDR corrected).

To test the second prediction, we again classified age group from EEG RDMs but shifted the time points for older participants backward by the observed peak latency delay of 21ms for image classification (Fig. 4c). We reasoned that if younger and older participants differ in timing but share a similar representational geometry, then age group classification should break down at the time point of younger adults’ peak classification (i.e., around 160ms). Considering the time window 100-300ms, we indeed observed that we can correctly classify age group before and after, but not at the peak latency of younger adults (Fig. 4d; 10,000 permutations, one-tailed permutation test, p<0.05, FDR corrected). This further strengthens the idea that temporal delays characterize the differences in the temporal dynamics of object processing in younger and older adults.

### 3.2 Aging delays object processing in high-level ventral visual cortex

**Figure 5.**
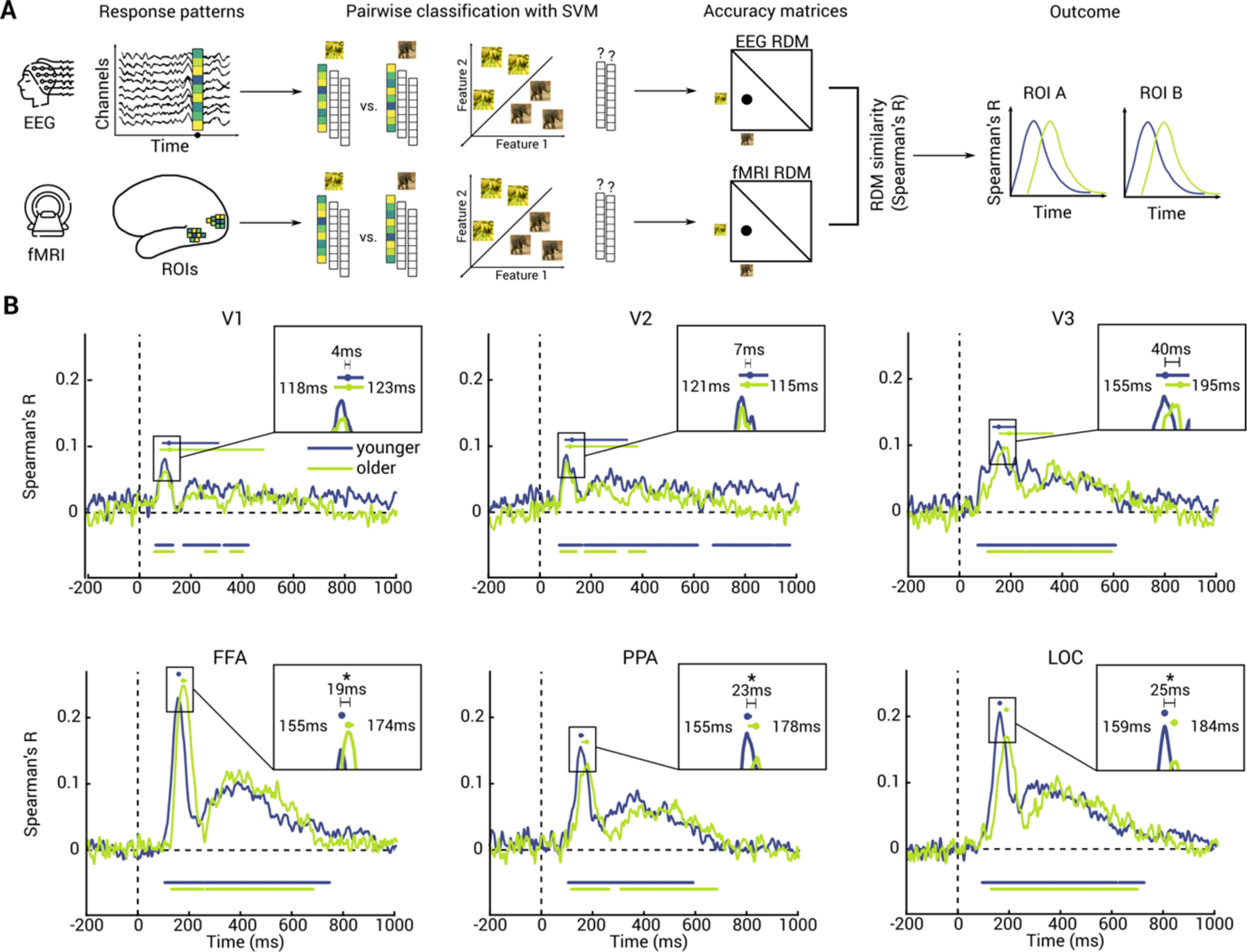
EEG-fMRI fusion. A) For EEG (top) and fMRI (bottom), we formed representational dissimilarity matrices (RDMs) from pairwise classification accuracies. We then correlated all possible combinations of RDMs using Spearman’s R, resulting in correlation time per ROI. B) Results for V1 (top left), V2 (top middle), V3 (top right), FFA (bottom left), PPA (bottom middle) and LOC (bottom right) for younger (blue) and older (green) adults. Shaded areas around curves indicate standard error of the mean. Significance time points are indicated below curves (10,000 permutations, one-tailed permutation test, p<0.05, cluster threshold q<0.05). Dots above curves indicate peaks of the fusion curves and error bars represent the 95% CI. Insets indicate peak latency for younger adults on the left, peak latency for older adults on the right and peak latency differences on the top. Stars indicate significant peak latency differences between age groups (p < 0.05; bootstrap test with 10,000 bootstraps).

To relate the observed temporal dynamics to cortical sources measured with fMRI, we used representational M/EEG-fMRI fusion ^24,27^. For this, we assessed response patterns across the processing hierarchy of the ventral visual stream, i.e., the “what pathway” of visual processing. Our analysis included six region-of-interest (ROI): early-to mid-level areas V1, V2, V3 and high-level areas fusiform face area (FFA) ^28^, parahippocampal place area (PPA) ^29^ and lateral occipital complex (LOC)30.

We again used time-resolved image classification results as EEG RDMs, and equivalently calculated ROI-based RDMs by classifying images from fMRI activation patterns. We then correlated ROI RDMs to time-resolved EEG RDMs for every participant, yielding a time course of representational similarity for each ROI (Fig. 5a).

We found representational correspondence between EEG and fMRI signals for all ROIs and both age groups (Fig. 5b; 10,000 permutations, one-tailed permutation test, p<0.05, cluster threshold q<0.05) with comparable peak accuracies and curve shapes. This extends previous research on mapping visual object recognition in space and time from younger to older participants and provides a firm basis for direct comparison across age groups.

To determine the regions at which processing delays occur we compared peak latencies for younger and older adults in each ROI (Fig. 5b insets; 10,000 bootstraps, bootstrap test against 0, p<0.05). We found significant peak delays in older compared to younger adults in FFA, PPA and LOC, but not V1, V2 and V3.

This results pattern was qualitatively consistent across analysis choices, indicating the robustness of the results (see Fig. 5b and Supplementary Table 2 for subject-specific analyses; Supplementary Figure 1 and Supplementary Table 3 for correlating subject-wise fMRI RDMs to average EEG RDMs; Supplementary Figure 2 and Supplementary Table 4 for correlating subject-wise EEG RDMs to average fMRI RDMs).

This clarifies that age-related processing delays in visual object processing occur due to changes in processing in high-level rather than low-to mid-level ventral visual cortex.

### 3.3 Object representations decrease in distinctiveness in older adults

**Figure 6.**
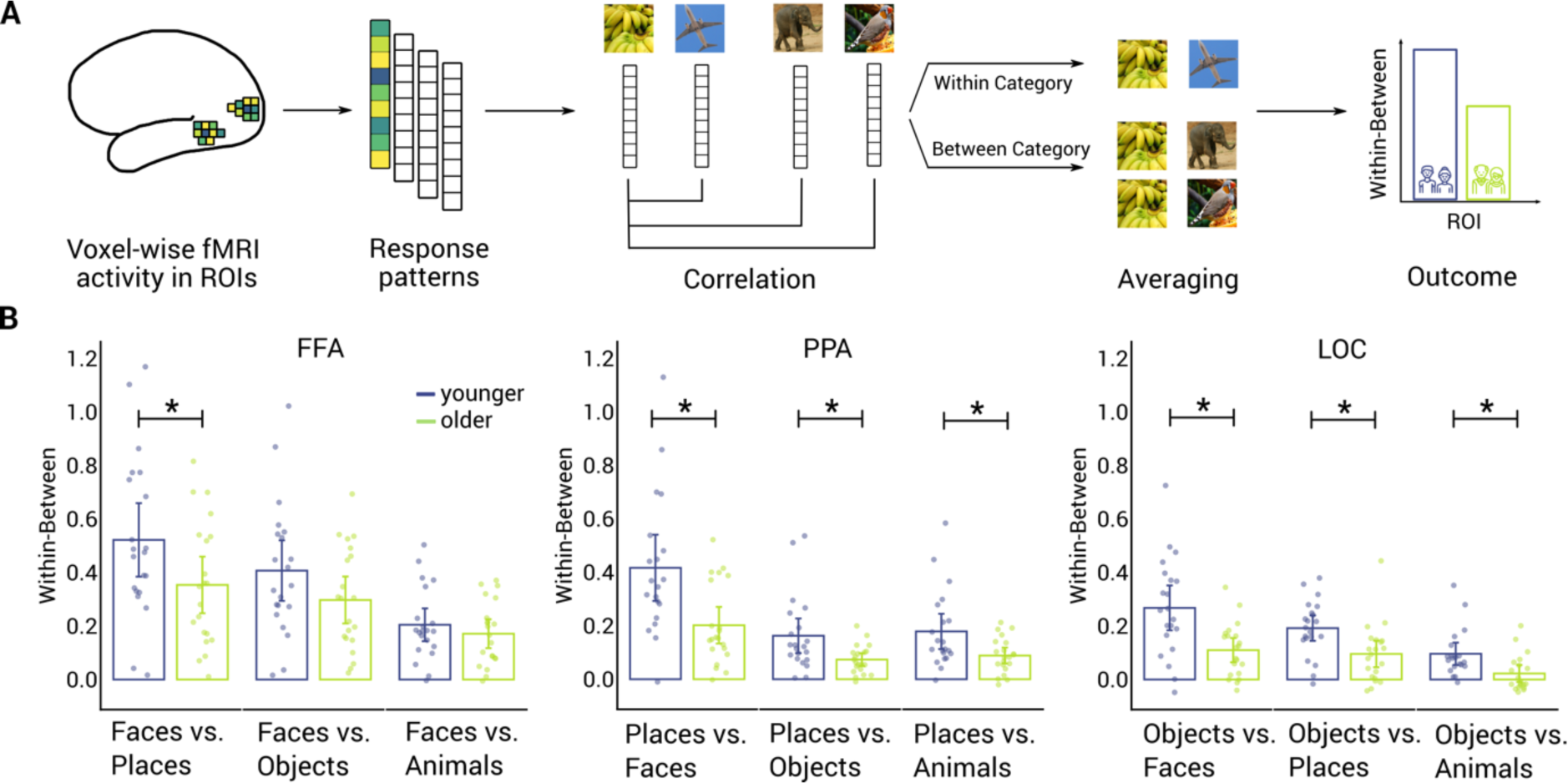
Pattern similarity analysis. A) We extracted fMRI voxel patterns from regions of interest (ROIs). We then correlated the fMRI voxel patterns of a given image and all other images belonging to the same category and averaged them into a within-category score. We also correlated the fMRI voxel patterns of a given image and all other images belonging to different categories and averaged them into a between-category score. We calculated the difference of the within- and between-category scores and compared the difference scores between age groups in different ROIs. B) Results of the pattern similarity analysis in the fusiform face area (FFA, left), parahippocampal place area (PPA, middle) and lateral occipital cortex (LOC, right) for younger (blue) and older (green) adults (10,000 permutations, one-tailed permutation test, p<0.05, FDR corrected). Error bars represent 95% confidence intervals. Dots represent single subject data. Stars indicate significant age groups differences.

The analyses above indicated high-level ventral visual cortex as the locus of visual processing delays in older participants. This raises the question of the nature of age-related changes in high-level ventral visual cortex. Guided by previous aging research ^31–33^, we focused on the concept of neural distinctiveness that has been observed to decrease in older adults. That is, we determined the degree to which object representations differed for preferred versus non-preferred categories in high-level ventral visual cortex. For this, we assessed the three ROIs that demonstrated processing delays above and for which category preferences can be defined: the face-preferring FFA, the place-preferring PPA and the object-preferring LOC.

Specifically, as measure of neural distinctiveness, we calculated how distinct brain response patterns are within and between categories. For each participant and each category (i.e., faces, places, objects and animals), we correlated the fMRI voxel patterns of a given image and all other images belonging to the same category (see Fig. 6a). We then Fisher z-transformed all resulting correlations and averaged them into a within-category score. We repeated the same process for a between-category score, i.e., correlating fMRI voxel patterns of a given image and all other images belonging to different categories. Finally, we calculated the difference of the within-and between-category scores for all category combinations, yielding the distinctiveness score.

For each ROI, we calculated the distinctiveness score based on comparing the region’s preferred category to each non-preferred category (Fig. 6b; 10,000 permutations, one-tailed permutation test, p<0.05, FDR corrected). Overall, younger participants had significantly higher distinctiveness scores than older participants in all three regions. More specifically, in PPA and LOC, this result did not depend on the non-preferred category, but in FFA, we found a significant age group difference only when comparing faces to places.

This results pattern was qualitatively consistent across analysis choices, indicating the robustness of the results (see Figure 6b as well as Supplementary Table 5 for results when using all 20 runs to maximize signal-to-noise ratio and Supplementary Figure 3 as well as Supplementary Table 6 for results when using the first 2 experimental runs to closely replicate previous studies^32,33^).

Together, these results indicate that object representations across high-level ventral visual cortex are less distinct in healthy older participants than in younger participants.

### 3.4 Visual object representations are related to behavior in younger and older participants

**Figure 7.**
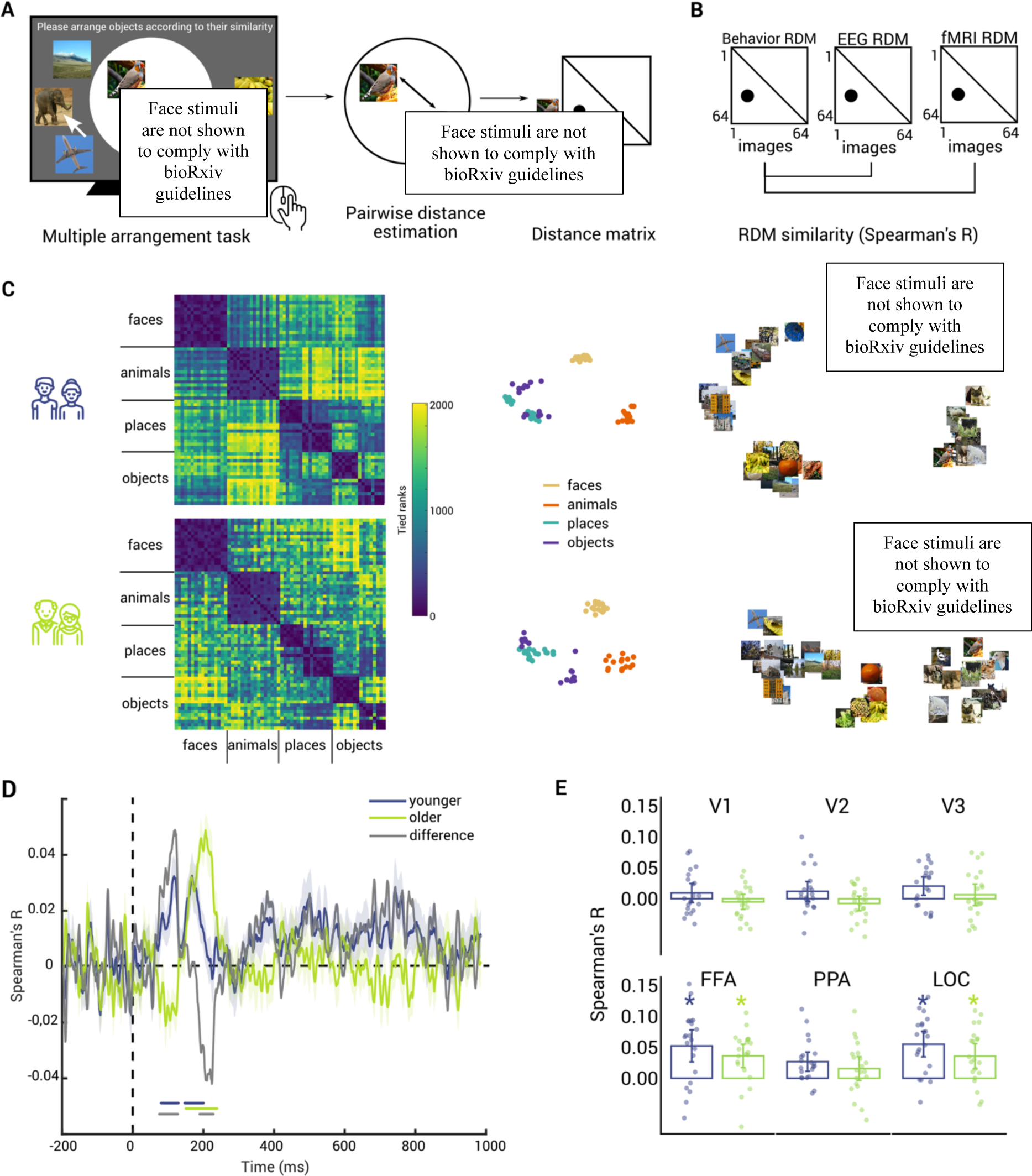
Behavioral measures and their representational similarity to neural measures. A) In the behavioral task, participants arranged the stimuli according to their perceived similarity on a computer screen via drag-and-drop by mouse. We then formed behavioral RDMs from distances between each pair of images and correlated them to time-point wise EEG RDMs and ROI-wise fMRI RDMs using Spearman’s R. B) Structure of the behavioral RDMs of younger (top) and older (bottom) adults revealed by tied ranks (left) or the first two dimensions of multidimensional scaling (middle and right). D) Results of behavior-EEG RSA for younger adults (blue), older adults (green) and age group difference (grey). Shaded areas around curves indicate standard error of the mean. Significance time points are indicated below curves (10,000 permutations, one-tailed permutation test, p<0.05, cluster threshold q<0.05). E) Results of behavior-fMRI RSA in V1 (top left), V2 (top middle), V3 (top right), FFA (bottom left), PPA (bottom middle) and LOC (bottom right) for younger (blue) and older (green) adults (10,000 permutations, one-tailed permutation test, p<0.05, FDR corrected). Error bars represent 95% confidence intervals. Dots represent single subject data. Colored stars indicate significant effects per age group.

So far, we identified representations and age-related changes therein by establishing statistical dependencies between brain signals and the stimulus material from the experimenter’s perspective. However, this does not distinguish whether the representations identified are used by the brain itself for visual behavior, or whether they are epiphenomenal ^34–36^.

As a first step towards arbitrating between these hypotheses, we assessed the perceived similarity of image stimuli in our data sets. The rationale is that if two objects are perceived to be similar, they will elicit similar behavior, e.g., when identifying novel objects and for categorization ^37–40^. Thus, a relationship between neural representations and perceived similarity ratings indicates that the identified representations are suitably formatted to guide visual behavior ^41–43^.

We assessed perceived similarity in younger and older adults between all images simultaneously using an iterative, computerized multiple arrangement task ^17,44^. We asked participants to arrange the 64 stimulus images on a screen via drag and drop with a mouse according to their similarity, with similar images located closer together and dissimilar images located further apart (Fig. 7a). This yielded behavioral RDMs for younger and older participants based on the distances between images.

We found that younger and older adults perceive the image set similarly. In both age groups, a visual inspection of the raw behavioral RDMs (Fig. 7b left) revealed a pronounced category structure, differentiating between faces, animals, places and objects. A two-dimensional projection of the high-dimensional RDM that preserved distances between images using multidimensional scaling confirmed this observation qualitatively (Fig. 7b right). We also compared the correlation of behavioral RDMs between age groups to an empirical distribution of RDM correlations based on 10,000 random group splits ignoring age group information. The result indicated that the correlation between younger and older adults falls within the central 95% of the distribution (p=0.2453 for left tail), providing no evidence for a difference in RDMs based on age group. This indicates that younger and older adults perceive objects similarly and warrants further investigation on a strong empirical basis.

In a next step, we related behavioral RDMs to EEG and fMRI data using RSA for each age group separately (Fig. 7d).

Considering the temporal dynamics with EEG first, we observed a differential pattern for younger and older adults (Fig. 7e, 10,000 permutations, one-tailed permutation test, p<0.05, cluster threshold q<0.05). In younger adults, we observed significant effects in two early time windows, from 80-130ms and 148-203ms. In older adults the effects emerged relatively later and became significant in the time window 152-242ms. Contrasting the results for younger and older adults in a difference curve (Fig. 7e, gray, 10,000 permutations, two-tailed permutation test, p<0.05, cluster threshold q<0.05) revealed a stronger relationship for younger adults earlier in time (76-129ms) and a stronger relationship for older adults later in time (192-231ms). This shows that the visual object representations identified in both age groups using EEG are suitably formatted for behavior and suggests that they emerge later in older than in younger adults, consistent with the overall pattern of results.

The overall result pattern was qualitatively consistent across analysis choices, indicating the robustness of the results (see Fig. 7e for correlating subject-wise EEG RDMs to the averaged behavior RDM; Supplementary Figure 4a for correlating subject-wise behavior RDMs to average EEG RDMs; Supplementary Figure 4b for subject-specific analysis; see Supplementary Table 7 for statistical details of all analyses variants).

Finally, relating behavioral RDMs to ROIs across the ventral visual stream revealed significant effects in FFA and LOC for both younger and older adults (Fig. 7f; 10,000 permutations, one-tailed permutation test, p<0.05, FDR corrected). Age groups did not differ significantly in any ROI (all p>0.2185). This indicates that object representations identified in high-level, but not low- or mid-level visual cortex are suitably formatted to guide behavior in both age groups.

This results pattern was also qualitatively consistent across analysis choices, indicating the robustness of the results (see Fig. 7e and Supplementary Table 8 for correlating subject-wise ROI RDMs to the averaged behavior RDM; Supplementary Figure 5a and Table 9 for correlating subject-wise behavior RDMs to average ROI RDMs; Supplementary Figure 5b and Table 10 for subject-specific analysis).

## 4. Discussion

We assessed how brain aging impacts visual representations. Our experimental strategy combining EEG, fMRI and behavioral assessment in younger and older adults yielded four key findings: First, assessing the chronometry, we found that aging delays the formation of object representations. Second, relating chronometry to cortical loci, we found that age-related delays in the formation of object representations manifest in high-level rather than low-to mid-level ventral visual cortex. Third, we found that aging changes visual representations in these category-preferring high-level areas by reducing their neural differentiation. Fourth, we ascertained ecological relevance by showing that the identified neural representations in younger and older adults were suitably formatted to guide visual behavior.

### 4.1 Aging delays the formation of visual object representations

Using time-resolved multivariate analysis of EEG data ^13,45,46^, we showed that the formation of visual representations is delayed in healthy aging: older adults showed lower classification accuracies in an early time window of ∼60-160ms, and their peak latency was delayed by ∼16-30ms. Our results concur with previous aging studies demonstrating largest delays in processing speed in the N170 time window ^47–49^ and increased peak latencies for event-related potential (ERP) components P1 ^50–53^, N170 ^54,55^ and M170 ^56^. Harvesting the power of multivariate analyses ^10–13^, we extend previous studies in three ways. First, our results establish a delay at all investigated levels of abstraction, from single images to (supra-)categories. This strengthens the empirical evidence for a robust age-related delay, affecting all processing levels. Second, using temporal generalization analysis ^23,24^, we show the persistent dynamics contributing to the observed delay. Third, using representational similarity analysis (RSA) ^25^, we establish that the delay indexes the emergence of similar rather than different representations in younger and older adults. Taken together, our results demonstrate that visual representations in younger and older adults are similar but formed at different speeds, indicating a profound age-related change in chronometry.

Apart from the observed early age-related processing difference, we also found that older adults have significantly lower classification accuracies around 700-900ms after stimulus onset. This might reflect reduced sustained attention in older adults: previous studies linked late positive potentials in this time window to participants allocating attentional resources to visual stimuli ^21,22^ and reported this component to be diminished in older compared to younger adults ^57^.

The success of classifying age group from EEG data underlines the robustness of the identified age-related differences and has promising translational potential. New effective and affordable biomarkers of pathological aging ^58,59^ are needed due to the global rise in the aging population, the prevalence of dementia among them and its considerable burden on low- and middle-income countries. Collecting EEG data during natural image viewing could be such a biomarker as it is non-invasive, affordable, not bound to hospital settings, independent of language and educational background and cognitively less effortful than memory tasks.

### 4.2 Age-related processing delays manifest in high-level ventral visual cortex

Our results clarify the exact locus at which age-related processing delays manifest functionally: using RSA-based EEG-fMRI fusion, we showed delayed processing in high-level rather than low- or mid-level ventral visual cortex. This concurs with previous studies showing that age-related delays in visual processing are not due to reductions in low-level vision such as visual acuity ^60^ or pupil size ^49^. It is further consistent with the idea that the ERP components P1 and N170, which are temporally aligned with the classification peaks reported above, have putative sources in high-level ventral visual cortex^61–63^.

Locating age-related processing delays to high-level ventral visual cortex illustrates the importance of our findings for research on high-level cognition in aging and its translational applications. As high-level ventral visual cortex enables object recognition but also bridges to memory systems such as the perirhinal cortex, memory studies should account for age-related feed-forward delays at this processing stage ^64^. Further, activity in high-level visual cortex during memory encoding is associated with higher cognitive reserve against Alzheimer’s pathology, emphasizing the relevance of our findings for clinical applications ^65^.

Our results also suggest a functional counterpart to age-related, anatomical changes in the ventral visual cortex. Previous studies reported age-related grey matter decreases in the fusiform cortex while primary visual cortex volume remained stable ^66,67^. The grey matter in fusiform gyri did not only reduce in older age but showed a strong inverted-U shape across the lifespan, in line with the last-in first-out theory of visual cortical development ^67^.

### 4.3 Aging reduces the neural differentiation in category-selective brain areas

Our results showed reduced neural differentiation in healthy aging across the high-level ventral visual cortex regions FFA, LOC and PPA. This finding concurs with previous studies using an equivalent multivariate approach and reporting age-related neural dedifferentiation in scene-selective areas ^31–33,68–70^, but diverge in showing neural dedifferentiation in object-selective LOC and face-selective FFA, too.

Notably, we found this pattern both when closely replicating aforementioned studies ^32,33^ by using only the first two stimulus presentations and when averaging across all stimulus presentations to increase signal-to-noise ratio. In line with previous univariate studies also reporting age-related neural dedifferentiation for faces ^68,71–73^ and objects ^71^, our results point towards neural dedifferentiation in aging being a phenomenon affecting the entire high-level ventral visual cortex.

Previous research suggests the neurotransmitter GABA as the underlying neural mechanism: the ventral visual cortices of younger and older adults differ in GABA levels ^74^, and individual differences in GABA levels predict individual differences in neural distinctiveness in older adults ^74^. Age-related decreases in GABA could cause reduced lateral inhibition in neurons in high-level ventral visual cortex, which, in turn, could result in reduced distinctiveness ^75,76^.

We cautiously note that our findings showed the same trend but did not replicate statistically (p>0.0959) when using a support vector machine (SVM) rather than a correlation-based classifier (Supplementary Figure 6). While SVM classifiers depend on a small subset of voxel values to define the decision boundary, correlation classifiers weigh all voxels equally. Arguably, non-boundary defining vectors are affected by aging first, but more voxels could be affected with advancing age or the progression into pathological aging. Future studies are needed to test these ideas.

### 4.4 A relationship between age-related delays and neural dedifferentiation?

Our analyses independently established high-level ventral visual cortex as the locus of age-related processing delay manifestation and neural dedifferentiation. Combining both observations into a unifying mechanism, we suggest that healthy aging primarily impacts visual object recognition by changing the chronometry of the underlying visual processes and that these age-related delays might, at least partly, drive reductions in neural distinctiveness. The proposed connection is specific as temporal delays did not manifest in other regions, and younger and older adults otherwise show similar temporal dynamics: magnitudes, trajectory shapes, representational geometries, persistent and transient aspects are comparable across age groups.

Future studies could test this hypothesis by categorizing middle-aged adults ^77^ based on whether their EEG-based classification peaks align with those of younger or older adults. We predict middle-aged adults characterized by peak delays comparable to older adults to show neural dedifferentiation.

### 4.5 Neural representations assessed are related to visual behavior

We found that the assessed neural representations are suitably formatted to guide visual behavior. This ascertains the ecological relevance of age-related cortical changes in visual processing for behavior and warrants against the risk that the effects observed are epiphenomenal and visible to the experimenter only ^34–36^. We note that the relation between behavior and neural representations as established via perceived similarity judgments is indirect and likely differs for other tasks ^78^. However, our results make the prediction that, independent of task, younger and older adults will show similar loci but delayed emergence of representations related to behavior.

### 4.6 Limitations

We deliberately chose to assess a fairly homogenous sample of 60-73-year-old adults without cognitive deficits, acute medical issues and medication, or physical constraints prohibiting enrollment in neuroimaging studies. This enabled us to collect EEG, fMRI and behavioral data from all participants to provide a first nuanced view of object representations in healthy aging. However, we acknowledge that this limits the generalizability of our findings to the general aging population. We encourage future studies to assess more inclusive samples, accounting for diverse factors of participant variability ^79^ and a wider age range ^77^, to provide a more detailed picture of the continuous aging spectrum.

### 4.7 Conclusion

Our results provide a comprehensive account of the nature of visual representations in healthy aging, encompassing their exact chronometry, locus, mechanism and ecological relevance. We reveal the impact of aging on core visual processing and highlight the importance of factoring in perception-related delays when investigating high-level cognition in aging.

## 5. Methods

### 5.1 Participants

We conducted three separate experiments with 21 younger (19-33 years) and 22 older (60-73 years) participants (see Table 1): an EEG, an fMRI and a behavioral experiment. The fMRI data of one older participant were excluded from further analyses due to low task accuracy in the scanner. All participants reported normal or corrected-to-normal vision and had intact contrast sensitivity as measured by the Freiburg Visual Acuity Test (FrACT)^80^. In addition, we screened older participants for cognitive impairments using the Mini-Mental State Examination (MMSE)^81^. None of the participants had to be excluded based on a cut-off criterion for cognitive impairment, i.e., a score below 27/30 points. All experiments were approved by the ethics committee of the Department of Education and Psychology of the Freie Universität Berlin and were conducted in accordance with the Declaration of Helsinki. All participants provided informed consent prior to the studies and received a monetary reward or course credit for their participation.

### 5.2 Stimuli

In all experiments, we presented participants with 64 natural images. The stimulus set consisted of 16 images in each of the four categories included: faces, animals, places, objects (Fig. 1a). We chose face stimuli from FairFace^82^, place stimuli from Places365^83^ and both animal and object stimuli from MS COCO^84^. All images are available in the OSF repository https://osf.io/xeukw/.

### 5.3 Experimental procedures

#### 5.3.1 EEG experiment

The EEG experiment consisted of one session, divided into 8 blocks. During the EEG experiment, we presented images at the center of a grey screen, overlaid with a white fixation cross in the center. In regular trials, we presented images for 500ms with an ISI randomly varying between 400 and 500ms. We instructed participants to fixate on the central cross without blinking. Every 4-6 trials, we presented an image of a “smurf” and instructed participants to react by pressing a button and blinking. In these catch trials, we presented images for 500ms with an ISI randomly varying between 1400 and 1500ms to avoid contamination of movement on subsequent trials (Fig. 1b left). In total, one EEG recording session consisted of 2048 regular trials and 54 catch trials, amounting to a total of 2560 trials.

#### 5.3.2 fMRI experiment

The fMRI experiment consisted of two sessions. In the first fMRI session, we recorded a structural image (∼7 minutes), 10 runs of the main experiment (∼45 minutes), and a fieldmap (∼3 minutes). The total duration of the first session was 55 minutes excluding breaks. The second session included an additional 10 runs of the main experiment (∼45 minutes), two functional localizer runs (∼10 minutes), and another fieldmap (∼3 minutes), amounting to a total duration of 58 minutes excluding breaks.

##### 5.3.2.1. fMRI main experiment

The experimental setup was similar to that for EEG but adapted to the specifics of fMRI. We asked participants to fixate on the central cross at all times and respond to fixation cross color changes (white to red) in occasionally appearing catch trials with a button press. In each regular trial, we presented images in a 750ms on-off-on-off design, resulting in a total trial duration of 3s (Fig. 1b right). Each regular trial - belonging to one of the 64 experimental conditions - was repeated once per run (run duration: 4.5 minutes) in random order. Regular trials were interspersed every 3-6 trials with a catch trial.

##### 5.3.2.2 fMRI localizer experiment

To define regions-of-interest (ROIs) in early visual and ventral visual stream areas, we performed a separate functional localizer experiment. The localizer consisted of 35 cropped images per experimental condition: faces, houses, objects and scrambled objects. The localizer experiment consisted of two runs each lasting 272s each. Each run started with 16s of fixation and comprised 12 blocks of faces, houses, objects and scrambled objects on a grey background. Each stimulation block lasted for 16s with presentations of 20 different images (500ms on, 300ms off), including two one-back repetitions that participants were instructed to respond to with a button press. We presented stimulation blocks in random order and interspersed them with 4 blank fixation blocks.

#### 5.3.3 Behavioral experiment

In a separate behavioral experiment, participants performed a Multiple Arrangement Task ^17,44^ to judge the perceptual similarity of the images used in the neuroimaging experiments. In this task, participants arranged the stimuli on a computer screen according to their perceived similarity. Images were presented outside of a white circular arena and had to be arranged inside the area, using a computer mouse to drag and drop them. We instructed participants to locate more similar images closer together and dissimilar images further apart, without specific instructions regarding the similarity definition. The first trial included all 64 stimuli; subsequent trials consisted of subsets. The subsets were chosen based on an adaptive procedure and, in general terms, included images that were arranged in close proximity in the initial trial (see ^44^ for details).

### 5.4 EEG data acquisition and preprocessing

We recorded EEG data using a 64 channel actiCap snap cap, a Brainvision actiCHamp amplifier and the Brainvision recorder software. In this system, the electrodes are placed according to the standard 10-10 system, with an additional ground and reference electrode on the scalp. We recorded data at a sampling rate of 1,000 Hz and online filtered between 0.03 and 100 Hz.

We preprocessed data offline using Fieldtrip in MATLAB 2021a. We epoched the data between -200 and 1000ms relative to stimulus onset and notch filtered at 50Hz. In addition, we performed baseline correction by subtracting the mean of the 200ms pre-stimulus interval from the entire epoch.

### 5.5 fMRI data acquisition, preprocessing and preparation

#### 5.5.1 fMRI acquisition

We acquired MRI data on a 3-T Siemens Tim Trio scanner with a 32-channel head coil. We obtained structural images using a T1-weighted sequence (1 mm^3^ voxel size). For the main experiment and the localizer run, we obtained functional images covering the entire brain using a multiband 4 sequence (multiband factor 4, TR=750ms, TE=30ms, flip angle=55°, 3 mm^3^ voxel size, 40 slices, slice thickness =3mm, FOV=222mm, matrix size=74×74, interleaved acquisition).

#### 5.5.2 fMRI preprocessing

We organized MRI data according to the Brain Imaging Data Structure (BIDS)^85^ using heudiconv^86^ and ReproIn^87^ and preprocessed them using fMRIPrep (version 1.4.0)^88^ using MNI152Lin as the output space. We did not apply surface preprocessing. We carried out all further fMRI analyses in MATLAB R2021a (www.mathworks.com).

#### 5.5.3 Univariate fMRI analysis

For all univariate fMRI analyses, we used SPM12 (http://www.fil.ion.ucl.ac.uk/spm/) and custom written code. For the main experiment, we modelled the fMRI responses to the 64 experimental conditions for each run using a general linear model (GLM). The onsets and durations of each image presentation entered the GLM as regressors and were convolved with a hemodynamic response function (hrf). Six movement parameters (pitch, yaw, roll, x-, y-, z-translation) as well as their first- and second-order derivatives, the six highest aCompCor noise components ^89^ and the corresponding discrete cosine-basis regressors entered the GLM as nuisance regressors. In total, the GLM included 64 regressors for the experimental conditions and 33 nuisance regressors in each run.

We repeated this GLM approach 20 times, using a different hrf for the convolution in each iteration. We obtained the hrfs from an openly available library (https://github.com/kendrickkay/GLMsingle), derived from the Natural Scenes dataset^90^. For each voxel, we then identified the hrf that yielded the minimum mean residual and extracted the beta values for the 64 image regressors from the according GLM. Compared to using a canonical hrf, this approach allows a better estimation of the true hrf as hrfs are variable across both participants, brain regions and age ^91,92^. Overall, the procedure resulted in 64 beta maps per run and participant.

For the localizer experiment, we modelled the fMRI response to four experimental conditions (faces, houses, objects, scrambled objects) entering block onsets and durations as regressors of interest and six movement parameters as nuisance regressors before convolving with the hrf. We generated four contrasts from the resulting beta parameter estimates. We localized activations by using the following contrasts: early visual cortex (EVC) by visual stimulation > baseline, fusiform face area (FFA) by faces > houses, parahippocampal place area (PPA) by houses > objects, lateral occipital complex (LOC) by objects > scrambled objects. In sum, this resulted in four t maps per localizer run per participant.

#### 5.5.4 Definition of regions-of-interest

We defined regions of interest (ROIs) in ventral visual stream by using a functional localizer and restricting results to anatomical masks. For V1, V2, and V3, we used masks from the probabilistic Wang atlas ^93^. For FFA, PPA, and LOC, we used parcels provided by the Kanwisher lab ^94^. First, we calculated the overlap between a localizer t map and the according mask. We then extracted the 300 highest voxels from the resulting overlap.

### 5.6 Classification from EEG data

To determine the amount of image, category or supra-category information present in brain measurements, we used a multivariate classification scheme ^10–13^. We either trained and tested classifiers on 64 image identities, 4 categories or 2 supra-categories. All classification analyses were carried out in MATLAB R2021a (www.mathworks.com) and relied on binary c-support vector classification (C-SVC) with a linear kernel as implemented in the libsvm toolbox (https://www.csie.ntu.edu.tw/cjlin/libsvm) ^95^. Furthermore, all analyses were conducted in a participant specific manner.

#### 5.6.1 Time-resolved EEG classification

To determine the timing with which object information emerges in the brain, we conducted time-resolved EEG classification ^13,45^. For each time point of the epoched EEG data, we extracted 64 EEG channel activations and arranged them into pattern vectors for each of the experimental conditions (64 for image, 4 for category and 2 for supra-category classification). All participants had 32 trials per image, totaling up to 2048 trials. To increase the SNR, we randomly assigned the trials into bins and averaged them into 8 new pseudo-trials. We then performed 8-fold leave-one-pseudo-run-out-cross validation. Specifically, we assigned seven pseudo-trials to the training set and then tested the SVM on the remaining, eighth pseudo-trial corresponding to data from the same two images or categories as in the training set but using held-out data for the testing set. This yielded percent classification accuracy (50% chance level) as output. Equivalent SVM training and testing was repeated for all combinations of image or category pairs.

For 64 images, this resulted in 2016 pairwise classification accuracies. For 4 categories, this resulted in 6 pairwise classification accuracies. For 2 supra-categories, this resulted in 1 classification accuracy. This procedure was repeated 100 times with random assignment of trials to pseudo-trials, and across all combinations of image or category pairs. We averaged results across condition pairs, folds, and iterations, yielding a classification accuracy time course reflecting how much object information was present at each time point in each participant.

#### 5.6.2 Time-resolved EEG searchlight in sensor space

We conducted an EEG searchlight analysis resolved in time and sensor space (i.e., across 64 EEG channels) to gain insights into which EEG channels contributed to the results of the time-resolved analysis described above. For the EEG searchlight, we conducted the time-resolved EEG classification as described above with the following difference: For each EEG channel ci, we conducted the same classification procedure and stored the classification accuracy at the position of c. After iterating across all channels and down-sampling the time points to a 5ms resolution, this yielded a classification accuracy map across all channels and time points in 5ms steps for each participant.

#### 5.6.3 Time generalization analysis of EEG data

To investigate the degree to which object representations generalized across points in time, we used the temporal generalization method ^13,23,24,45^.

For a given time point, we trained a classifier analogous to the time-resolved EEG classification procedure. Decisively, we then tested the classifier on pattern vectors from each time point in the epoch (downsampled to a 5ms resolution). We repeated this approach for every time point, yielding a two-dimensional matrix of classification accuracies that displayed the temporal dynamics across time.

#### 5.6.4 Representational similarity analysis of EEG RDMs

We compared the geometry of object representations across age groups using representational similarity analysis (RSA) ^25^. RSA characterizes the representational space of a measurement space (i.e., EEG data) with a representational dissimilarity matrix (RDM). RDMs aggregate pairwise distances between responses to all experimental conditions, thereby abstracting from the activity patterns of measurement units (i.e., EEG channels) to between-condition dissimilarities.

We used the classification results from the conducted image classification as a measure of (dis-)similarity relations between images. Classification accuracies can be interpreted as a measure of dissimilarity because two conditions have a higher classification accuracy when they are more dissimilar^24,26^. Thus, we assembled participant-specific RDMs for each EEG time point in the time course (downsampled to a 5ms resolution) from classification accuracies and extracted the lower triangular part of the respective RDMs (without the diagonal ^84^). We then averaged the participant-specific RDMs from 21 younger participants into one young-average RDM. In a final step, correlated (Spearman’s R) the participant-specific RDMs from 22 older participants with the young-average RDM for all time points combinations. This yielded a two-dimensional matrix of correlation coefficients indexed by time in younger adults on the x-axis and time in older adults on the y-axis. Accordingly, high correlations above the diagonal indicate that earlier time points in younger participants are correlated with later time points in older participants and vice versa. To directly compare correlation coefficients below and above the diagonal, we subtracted the lower triangular part of the matrix from the upper triangular part.

#### 5.6.5 Classifying age groups from EEG RDMs

For this analysis, we used the participant-specific and time point-wise EEG RDMs (downsampled to a 5ms resolution) described above. We trained an SVM to classify between the EEG RDMs of 20 younger and 20 older participants and tested it on the RDM of one left-out participant. We repeated this procedure for 43 folds and 1000 permutations at each EEG time point.

We first conducted the analysis in a time-resolved fashion using the same time-points for training and testing, yielding a time course of age group classification. To account for peak latency differences, we then repeated the classification analysis, keeping the same time points for younger participants but shifting the time points for older participants backward by the observed peak latency delay of 21ms. Importantly, this means that the resulting time course of age group classification represents time points for younger and time points plus delay for older adults.

### 5.7 RSA-based EEG-fMRI fusion

The rationale of the RSA-based EEG-fMRI fusion ^24,27^ was equivalent to the EEG RSA described above with the only difference being that we abstracted activity patterns from EEG channels and fMRI voxels to be able to compare representations across measurements spaces.

For EEG, we assembled participant-specific RDMs for each time point as described above. For fMRI, we performed a spatially resolved multivariate analysis, conceptually equivalent to the EEG image classification. Specifically, for each ROI, we extracted and arranged beta values into pattern vectors for each of the 64 images and 20 experimental runs. To increase the SNR, we randomly assigned run-wise pattern vectors into bins and averaged them into 4 pseudo-runs. We then performed 4-fold leave-one-pseudo-run-out-cross validation, training on 3 and testing on one pseudo-trial per classification iteration. We averaged classification accuracies across condition pairs and folds and organized them in participant-specific RDMs.

In a final step, we correlated (Spearman’s R) time-point wise EEG RDMs and ROI-wise fMRI RDMs, resulting in one time course per ROI. To ensure the robustness of our results, we conducted this analysis in three different variants: 1) correlating subject-wise EEG RDMs with subject-wise fMRI RDMs; 2) correlating subject-wise EEG RDMs with age-group averaged fMRI RDMs; 3) correlating age-group averaged EEG RDMs with subject-wise fMRI RDMs.

### 5.8 fMRI pattern similarity analysis

We employed a pattern similarity of fMRI data to investigate the neural distinctiveness of FFA, PPA and LOC. For each of these ROIs, we correlated the fMRI voxel patterns of a given image and all other images belonging to the same category. We repeated this for all 16 images from the same category, Fisher z-transformed all resulting correlations and averaged them into a within-category score. We repeated the same process for a between-category score, i.e., correlating fMRI voxel patterns of a given image and all other images belonging to different categories. Finally, we calculated the difference of the within- and between-category scores for all category combinations, yielding the distinctiveness score.

To ensure the robustness of our results, we conducted this analysis in two different variants. In the first variant, we used the average of all 20 runs to maximize signal-to-noise ratio. In the second variant, we only used data from the first two fMRI runs and the computed within- and between-category scores across trials of different runs to closely replicate the design of previous studies ^32,33^.

### 5.9 Analysis of behavioral measures

We constructed a similarity rating RDM from the multiple arrangement task. Each trial of the task provides a partial RDM. The partial RDMs were then combined into a full RDM following the algorithm described in the original publication ^44^. We analysed the correlation of behavioral RDMs across age groups by comparing it to an empirical distribution of RDM correlations based on 10000 random group splits ignoring age group information. We visualized the 64x64 behavioral RDMs using tied ranks and multidimensional scaling (MDS) ^96,97^. MDS is an unsupervised method enabling the visualization of similarity between objects in a distance matrix, i.e., it yielded coordinates in 2D space while preserving the distances between image pairs.

### 5.10 RSA of neural and behavioral measures

To determine the subset of neural image representations identified that is relevant for behavior ^14–17^, we compared perceptual similarity to image representations identified from EEG and fMRI signals using RSA ^25^.

We already described above how we constructed RDMs for EEG, fMRI, and behavior. We correlated the full behavioral RDMs with either time point-wise EEG or ROI-wise fMRI RDMs. For EEG, this analysis resulted in one correlation time course per participant. For fMRI, this resulted in one correlation value per participant and ROI.

To ensure the robustness of our results, we conducted this analysis in three different variants: 1) correlating age-group averaged behavior RDMs with subject-wise neural RDMs; 2) correlating age-group averaged neural RDMs with subject-wise behavior RDMs; 3) correlating subject-wise neural RDMs with subject-wise behavior RDMs.

### 5.11 Statistical testing

We applied non-parametric statistical tests to avoid assumptions about the distribution of the data. We used one sample permutation tests to assess patterns in younger and older participants separately. We applied independent samples permutation tests to compare age groups directly. Tests were either right- or two-tailed and are indicated for each result separately.

#### 5.11.1 Permutation tests

We performed non-parametric permutation tests to test for time points in the EEG classification, EEG time-generalization matrix, EEG RSA, and EEG age group classification. We also used permutations tests for time points and channels in the EEG searchlight, for ROI and time courses in the RSA and ROIs in the pattern similarity analysis.

For each one-sample test, the null hypothesis was that the observed parameter (i.e., classification accuracy, correlation) came from a distribution with a median of chance level performance (i.e., 50% for pairwise classification; 0 correlation). We permuted the labels of the data for only younger or only older participants, i.e., performed a sign permutation test that randomly multiplies participant-specific data with +1 or -1. For each independent samples test, the null hypothesis was that median in older equals the median in younger participants. In this case, we permuted the group labels for younger and older participants.

For each of the 10,000 permutation samples, we computed the statistic of interest, which resulted in an empirical distribution of the data. We then converted the original statistic into p values maps and thresholded them at q < .05 to define supra-threshold clusters. In addition, we constructed an empirical distribution of maximum cluster size from the permutation samples and only report clusters with sizes thresholded at p<.05.

For the age group classification, pattern similarity analysis, and fMRI-behavior RSA, we corrected the resulting *P*-values for multiple comparisons using the false discovery rate (FDR) ^98^ at 5% level instead of a cluster-level correction.

#### 5.11.2 Bootstrap tests

We used bootstrapping to compute the peaks of EEG time courses per age group, EEG peak latency differences between age groups and behavioral RDM correlations between age groups. In each case we sampled the participant pool 10,000 times with replacement and calculated the statistic of interest and 95% confidence interval.

For the EEG peak latency differences, we bootstrapped the latency difference between the peaks of younger and older participants’ classification time courses. This yielded an empirical distribution that could be compared to zero. To determine whether peak latency differences in the EEG time courses were significantly different from zero, we computed the proportion of values that were equal to or smaller than zero. For the behavioral RDM correlations, we bootstrapped two random participant groups and correlated the RDMs of both groups. We then compared the actual correlation of behavioral RDMs between age groups to the empirical distribution, computing the proportion of values that were equal to or smaller than the observed correlation.

### 5.12 Data availability

The raw EEG and fMRI data are available on OpenNeuro via https://openneuro.org/datasets/ds005363/ and https://openneuro.org/datasets/ds005374. The preprocessed data and RDMs for EEG, fMRI and behavior as well as the analysis results can be accessed on OSF via https://osf.io/xeukw/.

### 5.13 Code availability

The code used in this study is available on Github via https://github.com/marleenhaupt/ORHA/.

## Acknowledgements

We thank Zejin Lu for assistance with the stimulus set creation as well as Antoniya Boyanova and Mirjam Marx for help with data collection. We further thank all participants for taking part in our experiments and the HPC Service of ZEDAT, Freie Universität Berlin, for computing time ^1^.

## Author contributions

M.H., D.D.G., and R.M.C. designed research. M.H. collected and analyzed data. M.H. and R.M.C. wrote the manuscript. D.D.G. and R.M.C acquired funding.

## References

1. Bennett, L., Melchers, B. & Proppe, B. Curta: A General-purpose High-Performance Computer at ZEDAT, Freie Universität Berlin. (2020).

2. Salthouse, T. A. Trajectories of normal cognitive aging. Psychol. Aging 34, 17–24 (2019).

3. Harada, C. N., Natelson Love, M. C. & Triebel, K. L. Normal cognitive aging. Clin. Geriatr. Med. 29, 737–752 (2013).

4. 4. Schieber, F. Vision and Aging. in Handbook of the Psychology of Aging vol. 2 129–161 (Elsevier, 2006).

5. Bennett, P. J., Sekuler, R. & Sekuler, A. B. The effects of aging on motion detection and direction identification. Vision Res. 47, 799–809 (2007).

6. Billino, J., Bremmer, F. & Gegenfurtner, K. R. Differential aging of motion processing mechanisms: Evidence against general perceptual decline. Vision Res. 48, 1254–1261 (2008).

7. Springer, S. D. et al. Disturbances in primary visual processing as a function of healthy aging. Neuroimage 271, 120020 (2023).

8. Koen, J. D. & Rugg, M. D. Neural Dedifferentiation in the Aging Brain. Trends Cogn. Sci. 23, 547–559 (2019).

9. Pichot, R. E., Henreckson, D. J., Foley, M. F. & Koen, J. D. Neural noise is associated with age-related neural dedifferentiation. 1–33 (2022) doi:10.1101/2022.11.17.516990.

10. Cichy, R. M. et al. Probing principles of large-scale object representation: Category preference and location encoding. Hum. Brain Mapp. 34, 1636–1651 (2013).

11. Cichy, R. M., Chen, Y. & Haynes, J. D. Encoding the identity and location of objects in human LOC. Neuroimage 54, 2297–2307 (2011).

12. Carlson, T. A., Hogendoorn, H., Fonteijn, H. & Verstraten, F. A. Spatial coding and invariance in object-selective cortex. Cortex 47, 14–22 (2011).

13. Isik, L., Meyers, E. M., Leibo, J. Z. & Poggio, T. The dynamics of invariant object recognition in the human visual system. J. Neurophysiol. 111, 91–102 (2014).

14. Cichy, R. M., Kriegeskorte, N., Jozwik, K. M., van den Bosch, J. J. F. & Charest, I. The spatiotemporal neural dynamics underlying perceived similarity for real-world objects. Neuroimage 194, 12–24 (2019).

15. Bankson, B. B., Hebart, M. N., Groen, I. I. A. & Baker, C. I. The temporal evolution of conceptual object representations revealed through models of behavior, semantics and deep neural networks. Neuroimage 178, 172–182 (2018).

16. Mur, M. et al. Human object-similarity judgments reflect and transcend the primate-IT object representation. Front. Psychol. 4, 1–22 (2013).

17. Charest, I. et al. Unique semantic space in the brain of each beholder predicts perceived similarity. Proc. Natl. Acad. Sci. U. S. A. 111, 14565–14570 (2014).

18. Kay, K. N., Naselaris, T., Prenger, R. J. & Gallant, J. L. Identifying natural images from human brain activity. Nature 452, 352–355 (2008).

19. DiCarlo, J. J. & Cox, D. D. Untangling invariant object recognition. Trends Cogn. Sci. 11, 333–341 (2007).

20. Vanrullen, R. The power of the feed-forward sweep. 3, 167–176 (2007).

21. Hajcak, G., Macnamara, A. & Olvet, D. M. Event-related potentials, emotion, and emotion regulation: An integrative review. Dev. Neuropsychol. 35, 129–155 (2010).

22. Weinberg, A. & Hajcak, G. The late positive potential predicts subsequent interference with target processing. J. Cogn. Neurosci. 23, 2994–3007 (2011).

23. King, J. R. & Dehaene, S. Characterizing the dynamics of mental representations: The temporal generalization method. Trends Cogn. Sci. 18, 203–210 (2014).

24. Cichy, R. M., Pantazis, D. & Oliva, A. Resolving human object recognition in space and time. Nat. Neurosci. 17, 455–462 (2014).

25. Kriegeskorte, N., Mur, M. & Bandettini, P. Representational similarity analysis - connecting the branches of systems neuroscience. Front. Syst. Neurosci. 2, 1–28 (2008).

26. Guggenmos, M., Sterzer, P. & Cichy, R. M. Multivariate pattern analysis for MEG: A comparison of dissimilarity measures. Neuroimage 173, 434–447 (2018).

27. Cichy, R. M. & Oliva, A. A M/EEG-fMRI Fusion Primer: Resolving Human Brain Responses in Space and Time. Neuron 107, 772–781 (2020).

28. Kanwisher, N., McDermott, J. & Chun, M. M. The Fusiform Face Area: A Module in Human Extrastriate Cortex Specialized for Face Perception. J. Neurosci. 17, 4302–4311 (1997).

29. Epstein, R. & Kanwisher, N. The parahippocampal place area: A cortical representation of the local visual environment. Neuroimage 7, 6–9 (1998).

30. Grill-Spector, K., Kourtzi, Z. & Kanwisher, N. The lateral occipital complex and its role in object recognition. Vision Res. 41, 1409–1422 (2001).

31. Koen, J. D., Hauck, N. & Rugg, M. D. The relationship between age, neural differentiation, and memory performance. J. Neurosci. 39, 149–162 (2019).

32. Srokova, S., Hill, P. F., Koen, J. D., King, D. R. & Rugg, M. D. Neural Differentiation is Moderated by Age in Scene-Selective, But Not Face-Selective, Cortical Regions. eneuro 7, ENEURO.0142-20.2020 (2020).

33. Srokova, S., Aktas, A. N. Z., Koen, J. D. & Rugg, M. D. Dissociative effects of age on neural differentiation at the category and item level. J. Neurosci. 2, JN-RM-0959-23 (2023).

34. de-Wit, L., Alexander, D., Ekroll, V. & Wagemans, J. Is neuroimaging measuring information in the brain? Psychon. Bull. Rev. 23, 1415–1428 (2016).

35. Reddy, L. & Kanwisher, N. Category Selectivity in the Ventral Visual Pathway Confers Robustness to Clutter and Diverted Attention. Curr. Biol. 17, 2067–2072 (2007).

36. Williams, M. A., Dang, S. & Kanwisher, N. G. Only some spatial patterns of fMRI response are read out in task performance. Nat. Neurosci. 10, 685–686 (2007).

37. Nosofsky, R. M. Choice, similarity, and the context theory of classification. *Journal of Experimental Psychology: Learning*, Memory, and Cognition vol. 10 104–114 (1984).

38. Ashby, F. G. & Perrin, N. A. Toward a unified theory of similarity and recognition. Psychol. Rev. 95, 124–150 (1988).

39. Shepard, R. N. Toward a Universal Law of Generalization for Psychological Science. Science *(80-.).* **237**, 1317–1323 (1987).

40. Edelman, S. Representation is representation of similarities. Behav. Brain Sci. 21, 449–498 (1998).

41. Grootswagers, T., Cichy, R. M. & Carlson, T. A. Finding decodable information that can be read out in behaviour. Neuroimage 179, 252–262 (2018).

42. Karapetian, A. et al. Empirically Identifying and Computationally Modelling the Brain– Behavior Relationship for Human Scene Categorization. J. Cogn. Neurosci. 1–19 (2023) doi:10.1162/jocn_a_02043.

43. Singer, J. J. D., Karapetian, A., Hebart, M. N. & Cichy, R. M. The link between visual representations and behavior in human scene perception. bioRxiv (2023) doi:10.1101/2023.08.17.553708.

44. Kriegeskorte, N. & Mur, M. Inverse MDS: Inferring dissimilarity structure from multiple item arrangements. Front. Psychol. 3, 1–13 (2012).

45. Carlson, T. A., Hogendoorn, H., Kanai, R., Mesik, J. & Turret, J. High temporal resolution decoding of object position and category. J. Vis. 11, 1–17 (2011).

46. Carlson, T., Tovar, D. A., Alink, A. & Kriegeskorte, N. Representational dynamics of object vision: The first 1000 ms. J. Vis. 13, 1–19 (2013).

47. Rousselet, G. A. et al. Age-related delay in information accrual for faces: Evidence from a parametric, single-trial EEG approach. BMC Neurosci. 10, 114 (2009).

48. Rousselet, G. A. et al. Healthy aging delays scalp EEG sensitivity to noise in a face discrimination task. Front. Psychol. 1, 1–14 (2010).

49. Bieniek, M. M., Frei, L. S. & Rousselet, G. A. Early ERPs to faces: Aging, luminance, and individual differences. Front. Psychol. 4, (2013).

50. Tobimatsu, S. Aging and Pattern Visual Evoked Potentials. Optom. Vis. Sci. 72, (1995).

51. Kolev, V., Falkenstein, M. & Yordanova, J. Motor-response generation as a source of aging-related behavioural slowing in choice-reaction tasks. Neurobiol. Aging 27, 1719–1730 (2006).

52. Yordanova, J., Kolev, V., Hohnsbein, J. & Falkenstein, M. Sensorimotor slowing with ageing is mediated by a functional dysregulation of motor-generation processes: Evidence from high-resolution event-related potentials. Brain 127, 351–362 (2004).

53. Čeponiene, R., Westerfield, M., Torki, M. & Townsend, J. Modality-specificity of sensory aging in vision and audition: Evidence from event-related potentials. Brain Res. 1215, 53–68 (2008).

54. Wiese, H., Schweinberger, S. R. & Hansen, K. The age of the beholder: ERP evidence of an own-age bias in face memory. Neuropsychologia 46, 2973–2985 (2008).

55. Gazzaley, A. et al. Age-related top-down suppression deficit in the early stages of cortical visual memory processing. Proc. Natl. Acad. Sci. U. S. A. 105, 13122–13126 (2008).

56. Nakamura, A. et al. Age-related changes in brain neuromagnetic responses to face perception in humans. Neurosci. Lett. 312, 13–16 (2001).

57. Gaál, Z. A., Nagy, B., File, D. & Czigler, I. Older Adults Encode Task-Irrelevant Stimuli, but Can This Side-Effect be Useful to Them? Front. Hum. Neurosci. 14, 1–10 (2020).

58. Prince, M. J. et al. World Alzheimer Report 2015 - The Global Impact of Dementia: An analysis of prevalence, incidence, cost and trends. (Alzheimer’s Disease International, 2015).

59. Mattap, S. M. et al. The economic burden of dementia in low- and middle-income countries (LMICs): a systematic review. BMJ Glob. Heal. 7, 1–14 (2022).

60. Price, D. et al. Age-related delay in visual and auditory evoked responses is mediated by white-and grey-matter differences. Nat. Commun. 8, (2017).

61. Di Russo, F., Martínez, A., Sereno, M. I., Pitzalis, S. & Hillyard, S. A. Cortical sources of the early components of the visual evoked potential. Hum. Brain Mapp. 15, 95–111 (2002).

62. Watanabe, S., Kakigi, R. & Puce, A. The spatiotemporal dynamics of the face inversion effect: A magneto- and electro-encephalographic study. Neuroscience 116, 879–895 (2003).

63. Itier, R. J. & Taylor, M. J. Source analysis of the N170 to faces and objects. Neuroreport 15, 1261–1265 (2004).

64. Martin, C. B. & Barense, M. D. Perception and Memory in the Ventral Visual Stream and Medial Temporal Lobe. Annu. Rev. Vis. Sci. 9, 409–434 (2023).

65. Vockert, N., et al. Cognitive Reserve Against Alzheimer’s Pathology Is Linked to Brain Activity During Memory Formation. medRxiv (2023).

66. Raz, N. et al. Regional brain changes in aging healthy adults: General trends, individual differences and modifiers. Cereb. Cortex 15, 1676–1689 (2005).

67. Douaud, G. et al. A common brain network links development, aging, and vulnerability to disease. Proc. Natl. Acad. Sci. U. S. A. 111, 17648–17653 (2014).

68. Voss, M. W. et al. Dedifferentiation in the visual cortex: An fMRI investigation of individual differences in older adults. Brain Res. 1244, 121–131 (2008).

69. Zheng, L. et al. Reduced Fidelity of Neural Representation Underlies Episodic Memory Decline in Normal Aging. Cereb. Cortex 28, 2283–2296 (2018).

70. Carp, J., Park, J., Polk, T. A. & Park, D. C. Age differences in neural distinctiveness revealed by multi-voxel pattern analysis. Neuroimage 56, 736–743 (2011).

71. Park, D. C. et al. Aging reduces neural specialization in ventral visual cortex. Proc. Natl. Acad. Sci. U. S. A. 101, 13091–13095 (2004).

72. Park, J. et al. Neural broadening or neural attenuation? Investigating age-related dedifferentiation in the face network in a large lifespan sample. J. Neurosci. 32, 2154–2158 (2012).

73. Zebrowitz, L., Ward, N., Boshyan, J., Gutchess, A. & Hadjikhani, N. Dedifferentiated face processing in older adults is linked to lower resting state metabolic activity in fusiform face area. Brain Res. 1644, 22–31 (2016).

74. Chamberlain, J. D. et al. GABA levels in ventral visual cortex decline with age and are associated with neural distinctiveness. Neurobiol. Aging 102, 170–177 (2021).

75. Wang, Y., Fujita, I. & Murayama, Y. Neuronal mechanisms of selectivity for object features revealed by blocking inhibition in inferotemporal cortex. Nat. Neurosci. 3, 807–813 (2000).

76. Wang, Y., Fujita, I., Tamura, H. & Maruyama, Y. Contribution of GABAergic inhibition to receptive field structures of monkey inferior temporal neurons. Cereb. Cortex 12, 62–74 (2002).

77. Dohm-Hansen, S. et al. The ‘middle-aging’ brain. Trends Neurosci. 47, 259–272 (2024).

78. Hebart, M. N., Bankson, B. B., Harel, A., Baker, C. I. & Cichy, R. M. The representational dynamics of task and object processing in humans. Elife 7, 1–21 (2018).

79. Wig, G. S. et al. Participant diversity is necessary to advance brain aging research. Trends Cogn. Sci. 28, 92–96 (2024).

80. Bach, M. The Freiburg Visual Acuity Test-Variability unchanged by post-hoc re-analysis. Graefe’s Arch. Clin. Exp. Ophthalmol. 245, 965–971 (2006).

81. Folstein, M. F., Folstein, S. E., McHugh, P. R. & Ingles, J. ‘Mini-mental state’. A practical method for grading the cognitive state of patients for the clinician. J. Psychiatr. Res. 12, 189– 198 (1975).

82. Kärkkäinen, K. & Joo, J. FairFace: Face Attribute Dataset for Balanced Race, Gender, and Age. (2019).

83. López-Cifuentes, A., Escudero-Viñolo, M., Bescós, J. & García-Martín, Á. Semantic-aware scene recognition. Pattern Recognit. 102, (2020).

84. Lin, T. Y. et al. Microsoft COCO: Common objects in context. Lect. Notes Comput. Sci. (including Subser. Lect. Notes Artif. Intell. Lect. Notes Bioinformatics) 8693 LNCS, 740–755 (2014).

85. Gorgolewski, K. J. et al. The brain imaging data structure, a format for organizing and describing outputs of neuroimaging experiments. Sci. Data 3, 160044 (2016).

86. Halchenko, Y. O. et al. nipy/heudiconv: v1.1.6. (2024) doi:10.5281/zenodo.11497270.

87. Kennedy, D. N. et al. Everything matters: The repronim perspective on reproducible neuroimaging. Front. Neuroinform. 13, 0–9 (2019).

88. Esteban, O. et al. fMRIPrep: a robust preprocessing pipeline for functional MRI. Nat. Methods 16, 111–116 (2019).

89. Behzadi, Y., Restom, K., Liau, J. & Liu, T. T. A component based noise correction method (CompCor) for BOLD and perfusion based fMRI. Neuroimage 37, 90–101 (2007).

90. Allen, E. J. et al. A massive 7T fMRI dataset to bridge cognitive neuroscience and artificial intelligence. Nat. Neurosci. 25, 116–126 (2022).

91. Polimeni, J. R. & Lewis, L. D. Imaging faster neural dynamics with fast fMRI: A need for updated models of the hemodynamic response. Prog. Neurobiol. 207, 102174 (2021).

92. West, K. L. et al. BOLD hemodynamic response function changes significantly with healthy aging. Neuroimage 188, 198–207 (2019).

93. Wang, L., Mruczek, R. E. B., Arcaro, M. J. & Kastner, S. Probabilistic maps of visual topography in human cortex. Cereb. Cortex 25, 3911–3931 (2015).

94. Julian, J. B., Fedorenko, E., Webster, J. & Kanwisher, N. An algorithmic method for functionally defining regions of interest in the ventral visual pathway. Neuroimage 60, 2357– 2364 (2012).

95. Chang, C.-C. & Lin, C.-J. Libsvm: A library for support vector machines. ACM Trans. Intell. Syst. Technol. 2, 1–27 (2011).

96. Kruskal, J. & Wish, M. Multidimensional Scaling. (SAGE Publications, Inc., 1978). doi:10.4135/9781412985130.

97. Shepard, R. N. Multidimensional Scaling, Tree-Fitting, and Clustering. Science (80-.). 210, 390–398 (1980).

98. Benjamini, Y. & Hochberg, Y. Controlling the False Discovery Rate: A Practical and Powerful Approach to Multiple Testing. J. R. Stat. Soc. Ser. B 57, 289–300 (1995).

