## Supplementary Material for "Healthy aging delays and dedifferentiates high-level visual representations"

**Supplementary Tables**

Suppl. Table 1. Statistical details of EEG decoding within time related to Figure 2B…………..3

Suppl. Table 2. Statistical details of subject-wise EEG-fMRI fusion related to Figure 5B..........4

Suppl. Table 3. Statistical details of EEG-averaged EEG-fMRI fusion………………………...5

Suppl. Table 4. Statistical details of fMRI-averaged EEG-fMRI fusion………………………..6

Suppl. Table 5. Statistical details of pattern similarity analysis averaged across fMRI runs related to Figure 6B…………………………………………………………………………7

Suppl. Table 6. Statistical details of pattern similarity analysis based on first 2 fMRI runs.........8

Suppl. Table 7. Statistical details of Behavior-EEG RSA………………………………………9

Suppl. Table 8. Statistical details of behavior-averaged Behavior-fMRI RSA related to Figure 7E……………………………………………………………………………………..10

Suppl. Table 9. Statistical details of fMRI-averaged Behavior-fMRI RSA…………………...11

Suppl. Table 10. Statistical details of subject-wise Behavior-fMRI RSA……………………..12

**Supplementary Figures**

Suppl. Figure 1. EEG-fMRI fusion with averaged EEG RDMs………………………………13

Suppl. Figure 2. EEG-fMRI fusion with averaged fMRI RDMs……………………….………14

Suppl. Figure 3. Pattern similarity analysis based on first two fMRI runs…………….………..15

Suppl. Figure 4. Behavior-EEG RSA variants…………………………………….…………16

Suppl. Figure 5. Behavior-fMRI RSA variants……………………………………….………17

Suppl. Figure 6. Pattern similarity analysis equivalent based on SVM classification………..18

| Classification | Measure | Younger | Older | Difference |
|---|---|---|---|---|
| Image | Peak | 161 | 181 | 21 |
| | CI | 158-169 | 175-186 | 11-26 |
| | p | | | 6x10^-4 |
| | Sig. time points | 60-1000 | 69-1000 | 60-160 |
| Category | Peak | 187 | 203 | 16 |
| | CI | 182-194 | 193-213 | 2-30 |
| | p | | | 0.0139 |
| | Sig. time points | 64-1000 | 72-1000 | 69-186; 705-973 |
| Animacy | Peak | 164 | 194 | 30 |
| | CI | 162-168 | 171-210 | 7-47 |
| | p | | | 2x10^-4 |
| | Sig. time points | 65-1000 | 78-1000 | 76-162; 707-898 |

**Supplementary Table 1. Statistical details of EEG decoding within time related to Figure 2B.** Notes: All values except p values are reported in milliseconds; sig. = significant.

| Region of interest | Measure | Younger | Older | Difference |
|---|---|---|---|---|
| V1 | Peak | 118 | 123 | 4 |
| | CI | 91-343 | 82-488 | -236-381 |
| | p | | | 0.4854 |
| V2 | Peak | 121 | 115 | 7 |
| | CI | 91-313 | 94-386 | -213-277.5 |
| | p | | | 0.4752 |
| V3 | Peak | 155 | 195 | 40 |
| | CI | 130-219 | 156-366 | -46-216 |
| | p | | | 0.0569 |
| FFA | Peak | 155 | 174 | 19 |
| | CI | 149-165 | 164-188 | 5-34 |
| | p | | | 0.0061 |
| PPA | Peak | 155 | 178 | 23 |
| | CI | 149-167 | 156-185 | 0.5-32 |
| | p | | | 0.025 |
| LOC | Peak | 159 | 184 | 25 |
| | CI | 154-164 | 174-193 | 14-36 |
| | p | | | 0.0002 |

**Supplementary Table 2**. **Statistical details of subject-wise EEG-fMRI fusion related to Figure 5B.** Notes: All values except p values are reported in milliseconds; sig. = significant.

| ROI | Measure | Younger | Older | Difference |
| --- | --- | --- | --- | --- |
| V1 | Peak | 105 | 245 | 140 |
| | CI | 95-110 | 105-715 | -1-122 |
| | p | | | 0.3559 |
| V2 | Peak | 105 | 220 | 115 |
| | CI | 90-110 | 105-660 | -1-113 |
| | p | | | 0.1286 |
| V3 | Peak | 265 | 355 | 85 |
| | CI | 100-485 | 170-530 | -48-80 |
| | p | | | 0.3053 |
| FFA | Peak | 165 | 185 | 20 |
| | CI | 165-170 | 185-185 | 3-5 |
| | p | | | <1x10^-4 |
| PPA | Peak | 215 | 275 | 60 |
| | CI | 155-380 | 180-530 | -39-74 |
| | p | | | 0.1810 |
| LOC | Peak | 170 | 195 | 25 |
| | CI | 165-175 | 185-385 | 2-43 |
| | p | | | 1x10^-4 |

**Supplementary Table 3**. **Statistical details of EEG-averaged EEG-fMRI fusion.** Notes: All values except p values are reported in milliseconds (ms) and are downsampled to a 5ms resolution; sig. = significant.

| ROI | Measure | Younger | Older | Difference |
|---|---|---|---|---|
| V1 | Peak | 105 | 150 | 45 |
| | CI | 105-110 | 105-255 | -1-30 |
| | p | | | 0.3180 |
| V2 | Peak | 105 | 135 | 30 |
| | CI | 95-110 | 100-255 | -1-30 |
| | p | | | 0.1864 |
| V3 | Peak | 160 | 175 | 15 |
| | CI | 155-170 | 170-180 | 1-5 |
| | p | | | 0.0156 |
| FFA | Peak | 160 | 175 | 15 |
| | CI | 155-165 | 170-180 | 2-5 |
| | p | | | 4x10^-4 |
| PPA | Peak | 160 | 180 | 20 |
| | CI | 155-165 | 170-185 | 2-5 |
| | p | | | 5x10^-4 |
| LOC | Peak | 160 | 180 | 20 |
| | CI | 155-170 | 175-185 | 2-6 |
| | p | | | <1x10^-4 |

**Supplementary Table 4**. **Statistical details of fMRI-averaged EEG-fMRI fusion.** Notes: All values except p values are reported in milliseconds (ms) and are downsampled to a 5ms resolution; sig. = significant.

| Comparison | Mean Younger | Mean Older | p value<br>Age Difference |
|---|---|---|---|
| PPA: P vs. F | 0.4552 | 0.2019 | 8 x 10^-4 |
| PPA: P vs. A | 0.1788 | 0.0855 | 0.0050 |
| PPA: P vs. O | 0.1625 | 0.0734 | 0.0027 |
| LOC: O vs. F | 0.2676 | 0.1097 | 8 x 10^-4 |
| LOC: O vs. A | 0.0882 | 0.0183 | 0.0049 |
| LOC: O vs. P | 0.1919 | 0.0954 | 0.0028 |
| FFA: F vs. A | 0.2045 | 0.1711 | 0.1994 |
| FFA: F vs. P | 0.5218 | 0.3535 | 0.0238 |
| FFA: F vs. O | 0.4070 | 0.2972 | 0.0591 |

**Supplementary Table 5**. **Statistical details of pattern similarity analysis averaged across fMRI runs related to Figure 6B.** Notes: All values except p values are reported indicate within-between scores. A=animals, F=faces, O=objects, P=places; FFA= fusiform face area, LOC= lateral occipital complex, PPA= parahippocampal place area.

| Comparison | Mean Younger | Mean Older | p value<br>Age Difference |
|---|---|---|---|
| PPA: P vs. F | 0.0629 | 0.0205 | 6 x 10^-4 |
| PPA: P vs. A | 0.0325 | 0.0124 | 0.0025 |
| PPA: P vs. O | 0.0298 | 0.0080 | 4x 10^-4 |
| LOC: O vs. F | 0.0501 | 0.0045 | 1x 10^-4 |
| LOC: O vs. A | 0.0174 | -0.0066 | 0.0021 |
| LOC: O vs. P | 0.0349 | 0.0152 | 0.0232 |
| FFA: F vs. A | 0.0316 | 0.0318 | 0.5100 |
| FFA: F vs. P | 0.0897 | 0.0469 | 0.0048 |
| FFA: F vs. O | 0.0669 | 0.0416 | 0.0248 |

**Supplementary Table 6**. **Statistical details of pattern similarity analysis based on first 2 fMRI runs.** Notes: All values except p values are reported indicate within-between scores. A=animals, F=faces, O=objects, P=places; FFA= fusiform face area, LOC= lateral occipital complex, PPA= parahippocampal place area.

| Analysis variant | Measure | Younger | Older | Difference |
|---|---|---|---|---|
| Behavior avg. | Peak | 486 | 406 | -79 |
| | CI | 79-967 | 394-425 | -561-317 |
| | p | | | 0.6950 |
| EEG avg. | Peak | 947 | 404 | -543 |
| | CI | 708-974 | 69-753 | -901-(-217) |
| | p | | | 0.0074 |
| Subject-wise | Peak | 429 | 414 | -15 |
| | CI | 80-716.5 | 372.5-674 | -376-395.5 |
| | p | | | 0.7360 |

**Supplementary Table 7**. **Statistical details of Behavior-EEG RSA.** Notes: All values except p values are reported in milliseconds; avg=average. The analysis variant with averaged behavior relates to Figure 7D.

| ROI | Measure | Younger | Older | Difference |
|---|---|---|---|---|
| V1 | Spearman's R | 0.0093 | -0.0049 | |
| | p | 0.1354 | 0.8809 | 0.2185 |
| V2 | Spearman's R | 0.0120 | -0.0076 | |
| | p | 0.0849 | 0.9005 | 0.2185 |
| V3 | Spearman's R | 0.0205 | 0.0064 | |
| | p | 0.0088 | 0.3657 | 0.2185 |
| FFA | Spearman's R | 0.0523 | 0.0362 | |
| | p | 0.0015 | 0.0028 | 0.2747 |
| PPA | Spearman's R | 0.0269 | 0.0157 | |
| | p | 0.0019 | 0.1208 | 0.2747 |
| LOC | Spearman's R | 0.0551 | 0.0358 | |
| | p | 0.0011 | 0.0033 | 0.2185 |

**Supplementary Table 8**. **Statistical details of behavior-averaged Behavior-fMRI RSA related to Figure 7E.**

| ROI | Measure | Younger | Older | Difference |
| --- | --- | --- | --- | --- |
| V1 | Spearman’s R | 0.0108 | -0.0033 | |
| | p | 0.1273 | 0.6607 | 0.1531 |
| V2 | Spearman’s R | 0.0193 | 0.0002 | |
| | p | 0.0156 | 0.5422 | 0.1089 |
| V3 | Spearman’s R | 0.0471 | 0.0319 | |
| | p | 0.0002 | 0.0011 | 0.1271 |
| FFA | Spearman’s R | 0.0898 | 0.0551 | |
| | p | 0.0002 | 0.0002 | 0.0297 |
| PPA | Spearman’s R | 0.0549 | 0.0349 | |
| | p | 0.0002 | 0.0002 | 0.1026 |
| LOC | Spearman’s R | 0.0895 | 0.0628 | |
| | p | 0.0002 | 0.0002 | 0.0957 |

**Supplementary Table 9**. **Statistical details of fMRI-averaged Behavior-fMRI RSA.**

| ROI | Measure | Younger | Older | Difference |
|---|---|---|---|---|
| V1 | Spearman's R | 0.0071 | 0.0055 | |
| | p | 0.2092 | 0.2412 | 0.5951 |
| V2 | Spearman's R | 0.0066 | 0.0027 | |
| | P | 0.2092 | 0.3824 | 0.5951 |
| V3 | Spearman's R | 0.0201 | 0.0094 | |
| | p | 0.0198 | 0.1811 | 0.3507 |
| FFA | Spearman's R | 0.0469 | 0.0236 | |
| | p | 0.0044 | 0.0385 | 0.3507 |
| PPA | Spearman's R | 0.0215 | 0.0100 | |
| | P | 0.0115 | 0.2085 | 0.3507 |
| LOC | Spearman's R | 0.0443 | 0.0268 | |
| | p | 0.0044 | 0.0235 | 0.3507 |

**Supplementary Table 10**. **Statistical details of subject-wise Behavior-fMRI RSA.**

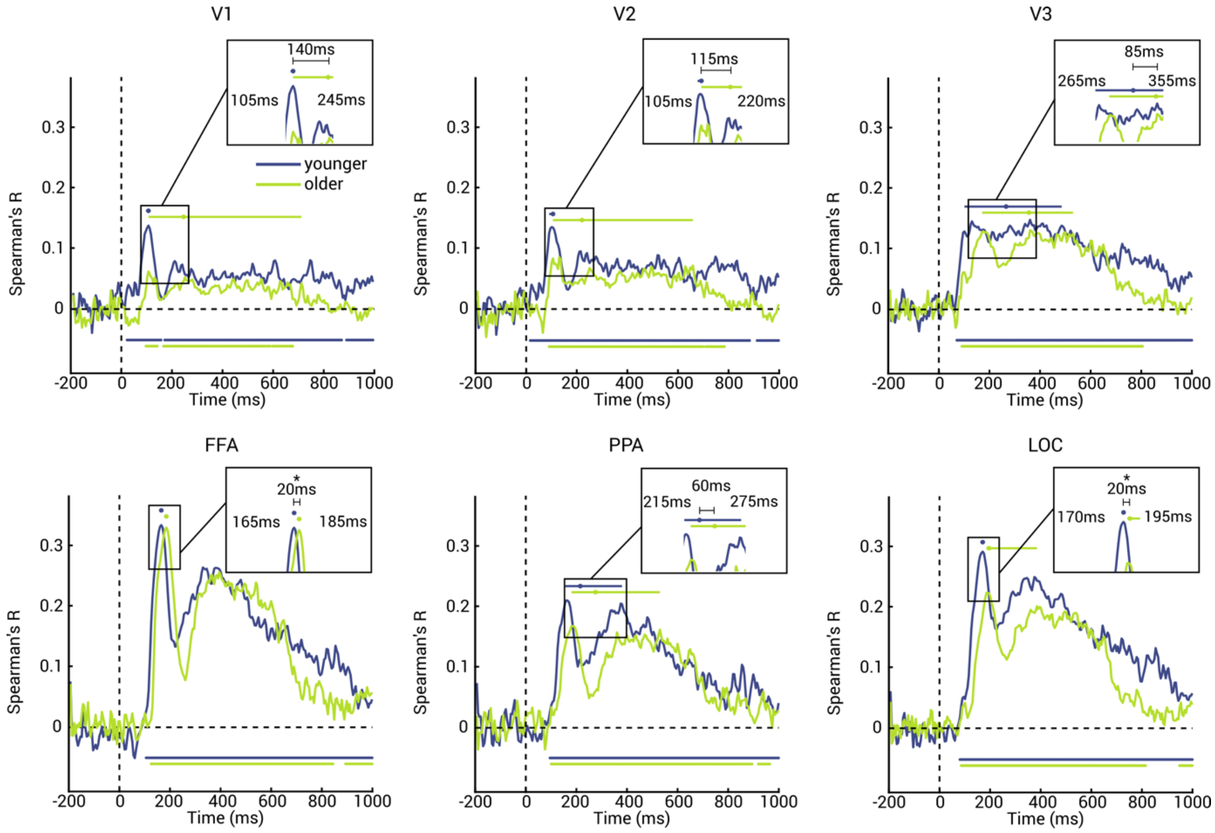

**Supplementary Figure 1. EEG-fMRI fusion with averaged EEG RDMs.** Results for V1 (top left), V2 (top middle), V3 (top right), FFA (bottom left), PPA (bottom middle) and LOC (bottom right) for younger (blue) and older (green) adults. Shaded areas around curves indicate standard error of the mean. Significance time points are indicated below curves (10,000 permutations, one-tailed permutation test, $p<0.05$, cluster threshold $q<0.05$). Dots above curves indicate peaks of the fusion curves and error bars represent the 95% CI. Stars indicate significant peak latency differences between age groups ($p < 0.05$; bootstrap test with 10,000 bootstraps).

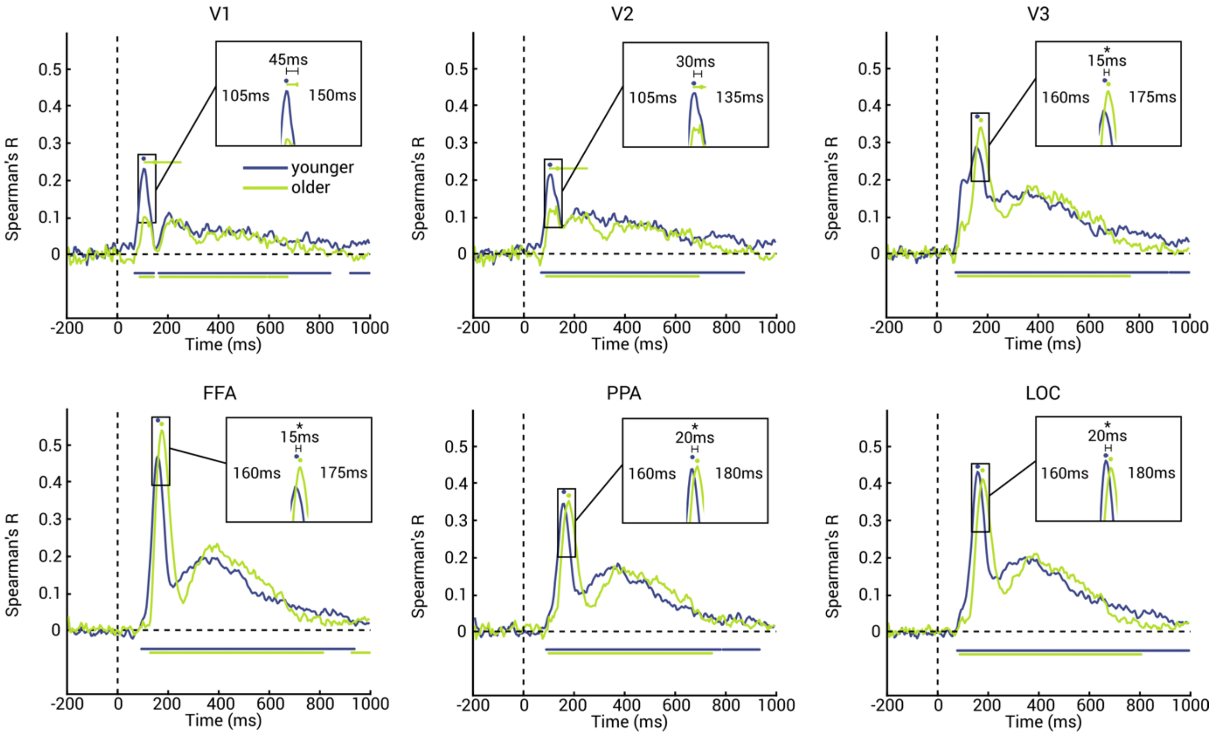

**Supplementary Figure 2. EEG-fMRI fusion with averaged fMRI RDMs.** Results for V1 (top left), V2 (top middle), V3 (top right), FFA (bottom left), PPA (bottom middle) and LOC (bottom right) for younger (blue) and older (green) adults. Shaded areas around curves indicate standard error of the mean. Significance time points are indicated below curves (10,000 permutations, one-tailed permutation test, $p<0.05$, cluster threshold $q<0.05$). Dots above curves indicate peaks of the fusion curves and error bars represent the 95% CI. Stars indicate significant peak latency differences between age groups ($p < 0.05$; bootstrap test with 10,000 bootstraps).

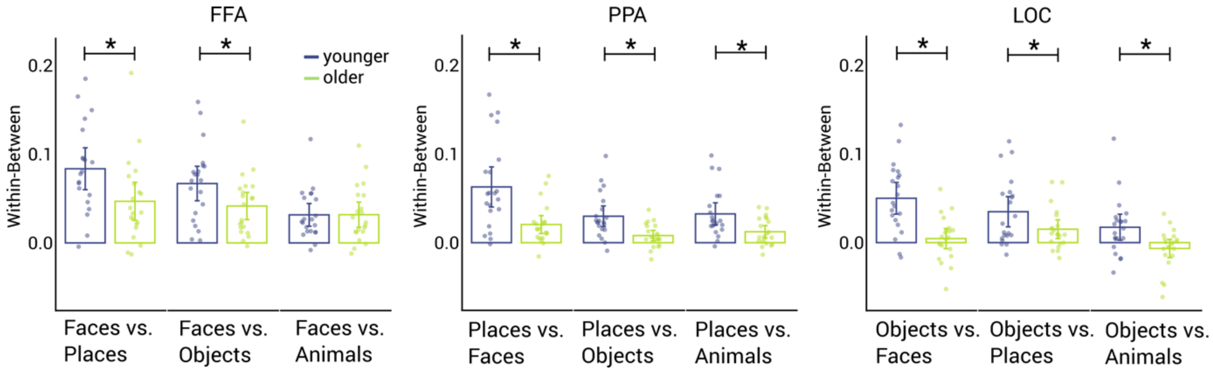

**Supplementary Figure 3. Pattern similarity analysis based on first two fMRI runs.** Results of the pattern similarity analysis in the fusiform face area (FFA, left), parahippocampal place area (PPA, middle) and lateral occipital cortex (LOC, right) for younger (blue) and older (green) adults (10,000 permutations, one-tailed permutation test, $p<0.05$, FDR corrected). Error bars represent 95% confidence intervals. Dots represent single subject data. Stars indicate significant age groups differences.

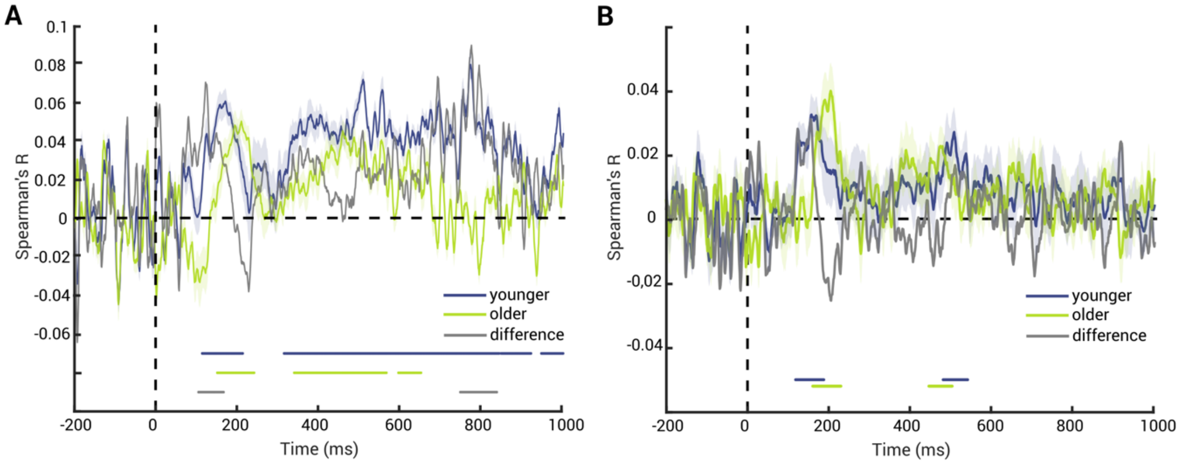

**Supplementary Figure 4. Behavior-EEG RSA variants.** Results for younger adults (blue), older adults (green) and age group difference (grey). Shaded areas around curves indicate standard error of the mean. Significance time points are indicated below curves (10,000 permutations, one-tailed permutation test, p<0.05, cluster threshold q<0.05). **(A)** Results for correlation averaged EEG RDMs with subject-wise behavior RDMs. **(B)** Results for correlating subject-wise EEG RDMs with subject-wise behavior RDMs.

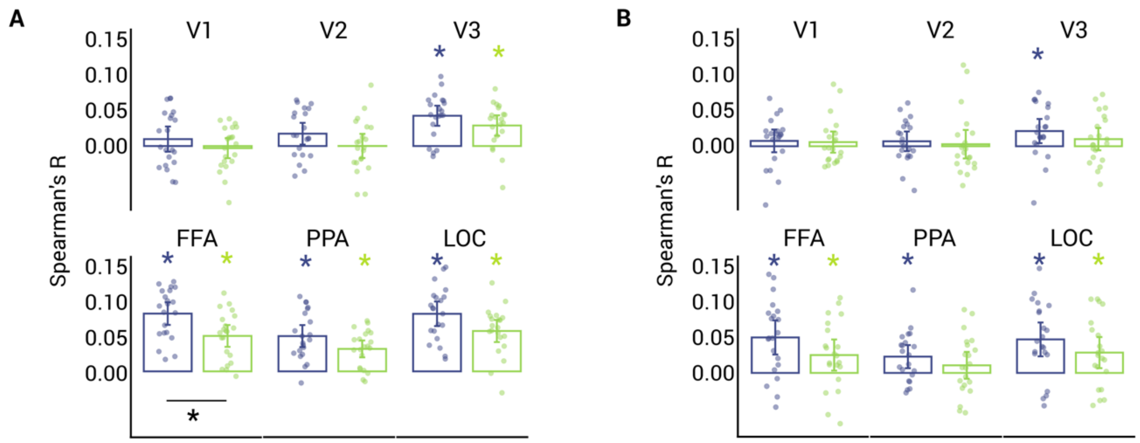

**Supplementary Figure 5. Behavior-fMRI RSA variants.** Results in V1 (top left), V2 (top middle), V3 (top right), FFA (bottom left), PPA (bottom middle) and LOC (bottom right) for younger (blue) and older (green) adults (10,000 permutations, one-tailed permutation test, $p<0.05$, FDR corrected). Error bars represent 95% confidence intervals. Dots represent single subject data. Stars indicate significant effects per age group. **(A)** Results for correlation averaged fMRI RDMs with subject-wise behavior RDMs. **(B)** Results for correlating subject-wise fMRI RDMs with subject-wise behavior RDMs.

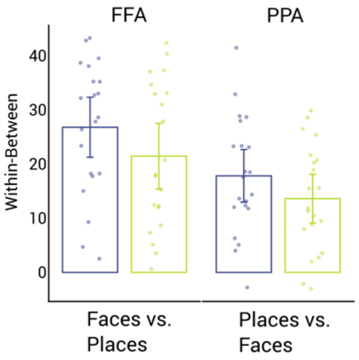

**Supplementary Figure 6. Pattern similarity analysis equivalent based on SVM classification.** Results of support vector machine classification in the fusiform face area (FFA) and parahippocampal place area (PPA) for younger (blue) and older (green) adults (10,000 permutations, one-tailed permutation test, $p<0.05$, FDR corrected). Error bars represent 95% confidence intervals. Dots represent single subject data. Stars indicate significant age groups differences.
